# Tissue tension permits β-catenin phosphorylation to drive mesoderm specification in human embryonic stem cells

**DOI:** 10.1101/2023.07.14.549074

**Authors:** Nadia M.E. Ayad, Johnathon N. Lakins, Ajinkya Ghagre, Allen J. Ehrlicher, Valerie M. Weaver

**Author notes:** Electronic correspondence.

## Abstract

The role of morphogenetic forces in cell fate specification is an area of intense interest. Our prior studies suggested that the development of high cell-cell tension in human embryonic stem cells (hESC) colonies permits the Src-mediated phosphorylation of junctional β-catenin that accelerates its release to potentiate Wnt-dependent signaling critical for initiating mesoderm specification. Using an ectopically expressed nonphosphorylatable mutant of β-catenin (Y654F), we now provide direct evidence that impeding tension-dependent Src-mediated β-catenin phosphorylation impedes the expression of Brachyury (T) and the epithelial-to-mesenchymal transition (EMT) necessary for mesoderm specification. Addition of exogenous Wnt3a or inhibiting GSK3β activity rescued mesoderm expression, emphasizing the importance of force dependent Wnt signaling in regulating mechanomorphogenesis. Our work provides a framework for understanding tension-dependent β-catenin/Wnt signaling in the self-organization of tissues during developmental processes including gastrulation.

## Introduction

The early embryo is remarkable for its shape-shifting capabilities. Extensive cell movement and tissue organization transform the inner cell mass of the embryo into three separate fates that ultimately form the body – ectoderm, endoderm, and mesoderm – through a process defined as gastrulation ^1^. Although the molecular pathways and motor proteins that drive morphogenetic movements during gastrulation have been extensively studied ^2–5^, how these forces impact the genetic determination of cell fate remains a topic of active investigation.

Cell fate occurs through a balance of diffusible secreted morphogens – in particular BMP, Nodal and Wnt ^6^ –, and their inhibitors ^7^ – such as Noggin, Lefty, DKK1 ^8^. However, emerging evidence suggest that mechanical forces driving and arising from morphogenetic movements also contribute to cell fate specification ^9–12^. β-catenin, a key developmental protein ^13, 14^, has been implicated in such mechanical signaling ^9, 10, 15–18^. Mechanical tension at adherens junctions (AJ) has been implicated in β-catenin phosphorylation of the C-terminus tyrosine 654 (Y654) in *Drosophila,* zebrafish, sea anemone and choanoflagellates ^19, 20^. This force-dependent β- catenin exposure is predicted to be permissive for Src-dependent Y654 phosphorylation, which reduces β-catenin binding to E-cadherin ^21^, leading to loss of AJ β-catenin ^22^, nuclear accumulation, and a Wnt-dependent mesodermal gene expression program ^23, 24^. Similarly, domains of high contractility in a two dimensional (2D) human gastruloid model have been shown to undergo Src dependent Y654 phosphorylation, accompanied by loss of AJ β-catenin, and robust mesodermal specification, thereby extending these findings to early mammalian development ^11^. Although collectively these developmental findings are compelling, we opted to directly interrogate the requirement of tension-dependent Y654 phosphorylation in vertebrate gastrulation.

To provide causal evidence for the role of force-mediated Src-dependent Y654 phosphorylation in mammalian gastrulation, we measured the effect of expressing a conserved nonphosphorylatable mutant of β-catenin (Y654F) in our human embryonic gastruloid model. Significantly, here we report direct evidence supporting a pivotal role for force-dependent Src-mediated β-catenin phosphorylation in human mesoderm specification. Evidence is presented to show that preventing Y654 phosphorylation had no impact on cell-generated contractility, Src activity or Src recruitment to the AJs but did compromise the ability of BMP4 to induce mesoderm differentiation, as demonstrated by a loss of epithelial-to-mesenchymal transition (EMT) and expression of the early mesodermal marker T (*TBXT* or *Brachyury*). Indeed, mesoderm signaling in the mutant cell line was rescued through addition of the β-catenin target Wnt3a, ^11^ and by increasing cytosolic β-catenin through inhibition of GSK3β using CHIR99021.

Our study provides definitive evidence for the evolutionary significance of the Y654 residue in mechanically regulating β-catenin activity during gastrulation. The findings provide important insight into the complex interplay between mechanical force and molecular signaling, not only advancing our understanding of embryonic development, but also tissue morphogenesis.

## Results

### Generation of mutant and wild-type cell lines

We first set about to generate hESCs that stably expressed a wild-type and a mutant nonphosphoratable β-catenin. To maximize any quantifiable phenotype and minimize levels of β-catenin overexpression we created stable recipient cell lines in which the endogenous β- catenin could be knocked down using an (orthogonal) IPTG inducible shRNAi lentiviral vector ^25^ (Sup.Fig1a). This vector constitutively expressed a (nuclear) mTurq2-NLS. Silent mutations in the shRNA recognition site of exogenously expressed β-catenin rendered it resistant to shRNA silencing.

Mindful of the potential effects of constitutive overexpression of β-catenin in driving spontaneous differentiation of hESCs we elected to use a doxycycline (DOX) inducible Sleeping Beauty transposon based vector ^26^ to generate stable cells lines that expressed the nonphosphorylatable Y654F mutant and wild type β-catenin as a control in an H9 hESC cell line with a CRISPR Cas9 generated T-mNeonGreen mesoderm reporter^11^. β-catenin was expressed as a polyprotein via a 2A peptide downstream of (nuclear) mScarlet-NLS (Sup.Fig1a).

To obtain a more homogenous undifferentiated population, we transiently induced mScarlet NLS T2A β-catenin using a brief 3h pulse of DOX followed by washout and flow sorting the following day, gating for expression of mScarlet and mTurq2 and absence of T-mNeonGreen (P7 gate on SupFig1b). Gating for high levels of mTurq2 correlated with high levels of IPTG inducible endogenous β-catenin knockdown. Gating for medium-high expression of mScarlet-NLS and negative T-mNeonGreen expression ensured that β-catenin levels would be sufficient for mesoderm signaling ^27^ but ideally not too high to induce differentiation and mesoderm signaling spontaneously when overexpressed ^28^. Accordingly, hESC pluripotency of these cell lines was retained, as validated by monitoring for expression of pluripotency markers *Nanog and Oct3/4* by quantitative polymerase chain reaction (qPCR) (SupFig1c). We did observe a modest but insignificant increase in condensed and fragmented nuclei (Fig.1f arrows) and in the level of cleaved-caspase-3 cells in the hESCs expressing the mutant β-catenin (SupFig1d-e). However, a Crystal Violet absorbance assay indicated cell proliferation was not significantly affected (SupFig1f). Β-catenin expression levels in the wild-type and mutant cell lines were similar to parental cell line levels (SupFig2a). Knockdown efficiency was measured by comparing β-catenin staining intensity levels (Sup.Fig2b and c) in both wild-type and mutant cell lines.

**Fig.1.**
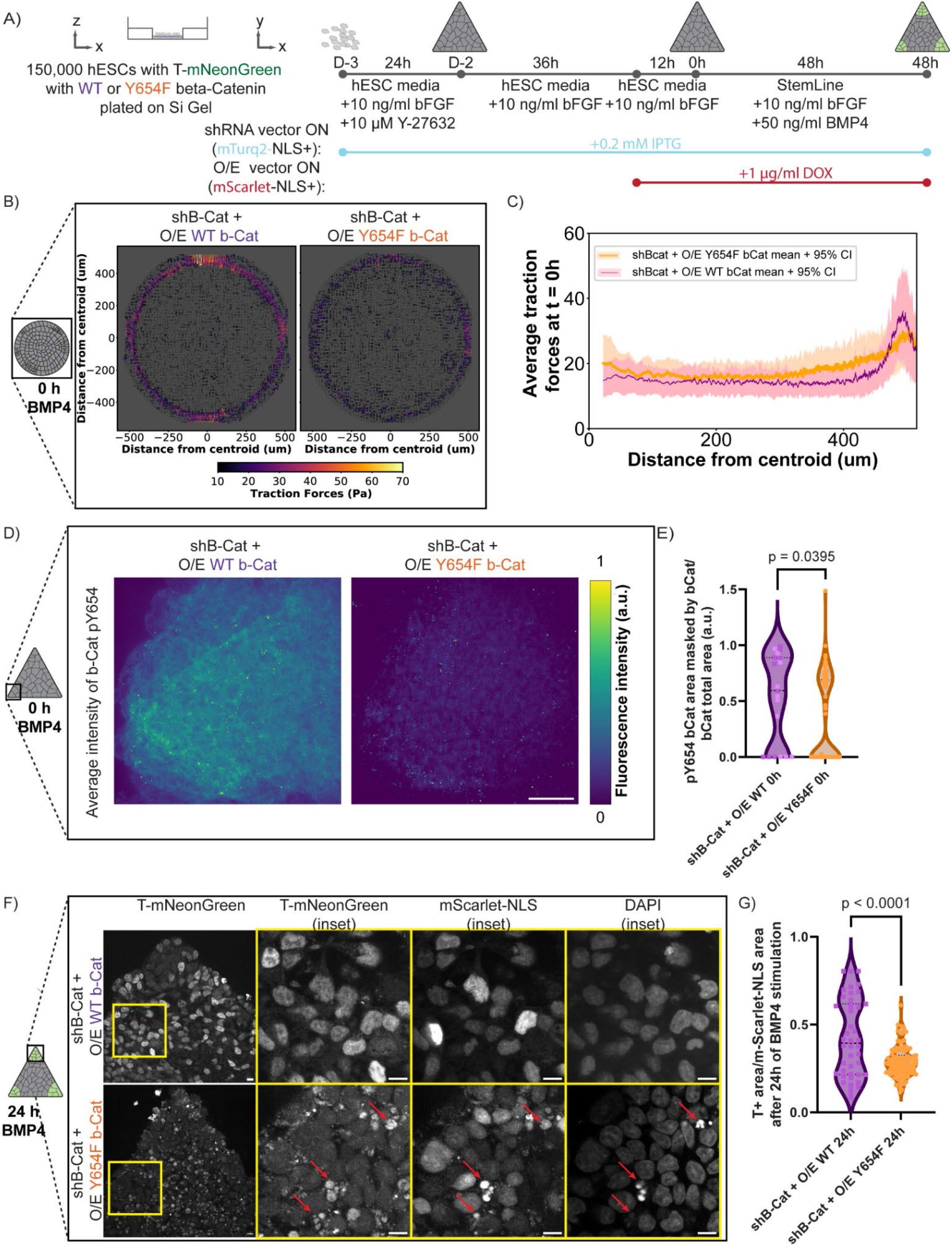
Phosphorylation of Y654 Β-catenin is mechanically activated and Y654F mutation prevents differentiation of T. A) Schematic of cell culture strategy for the differentiation of cells in this study. B) Average traction force distribution for wild-type and mutant circular colonies before the start of differentiation in circular colonies, with arrows indicating the direction and magnitude of the traction stress (color-coded for magnitude). N = 13 in Y654F and N = 10 for WT colonies for 2 independent experiments. C) Profile of average traction stresses for WT and Y654F circular colonies in D displaying mean (dark line) and 95% confidence intervals (shaded). D) Average intensity of phospho-Y654 β-catenin antibody in wild-type (WT) and mutant (Y654F) cells prior to the start of differentiation in the corner of triangle-shaped colonies. Scale bar = 10 μm E) Quantification of individual files of B. Images of phoshpho-Y654 β-catenin antibody were masked with the corresponding β-catenin image, then resulting area was normalized to the total β-catenin area. N= (8, 11) for WT and N = (35, 27, 31) for Y654F images. Unpaired two-tailed t-test was performed in data. F) Representative images of Y654F vs. WT cells after 24h of differentiation with BMP4, showcasing on the far left the expression of the T-mNeonGreen reporter. Inset shows a zoomed-in area of T-mNeonGreen (center-left), the corresponding expression of the mScarlet-NLS overexpressing vector (center-right) and the DAPI staining. Scale bar = 10 μm. G) Quantification of files shown in F. T+ area was thresholded with the StarDist algorithm, masked with the mScarlet-NLS StarDist-threshold and normalized to the total mScarlet-NLS area. N = (9, 22, 7) for WT and N= (65, 41) for Y654F cells. Unpaired two-tailed t-test was performed in data.

### Preventing Y654 phosphorylation of β-catenin impedes tension-stimulated mesoderm differentiation

We next plated our hESCs onto patterned gels of defined stiffness (Silicone-based gels of shear modulus; G’ ∼ 1.5kPa ^29^) that were conjugated with laminin-rich reconstituted basement membrane (rBM; Matrigel equivalent) ^30^ to generate either circular, square or triangular colonies. Cells were plated in hESC media containing 10 μM Rho-kinase inhibitor (Y27632; ROCKi) and 0.2 mM IPTG to induce β-catenin knockdown. After 24 hours, ROCKi was removed, the cells were washed, and fresh IPTG was added. 36 hours later, 1 μg/ml doxycycline was added to induce the ectopically expressed wild-type and mutant β-catenin vectors (Fig1a). After 12 hours of DOX induction of β-catenin transgene expression, which we defined as our initial timepoint (t = 0h), we began monitoring differentiation with the addition of 50 ng/ml BMP4 for up to 48 hours to induce a mesodermal program ^31^.

Traction Force Microscopy revealed that both wild type hESCs and our mutant hESC line developed high contractility at the periphery of circular colonies (Fig1d and e) ^11^, prior to the start of differentiation (shown here only on circles for visualization of profile quantification, but also present in the periphery of triangles and squares, as reported in ^11^).

Mutant and control hESCs were validated by staining for pY654 phosphorylation at the zero timepoint, focusing on regions within the patterned colonies with the highest contractility (Fig1b). Consistently, hESCs expressing the de-phospho-mimetic mutant β-catenin showed significantly less pY654 β-catenin in regions of high colony tension (shown here in triangle corners) as compared to the control hESCs expressing the wild-type β-catenin positive areas (Fig1c).

Monitoring for BMP sensitivity ^32^ by assessing nuclear pSMAD1/5 levels following 2 hours of BMP4 stimulation (50 ng/ml) revealed intact bimodal, edge-restricted nuclear pSMAD1/5, quantified in the circular colonies as the profile of nuclear signal from the centroid of the colony to its edge, as has been previously shown in dense 2D gastruloids (SupFig3a and b) ^33^. Importantly, wild-type colonies exhibited enhanced sensitivity to BMP4, as indicated by higher levels of nuclear p-SMAD/5 in the hESCs within the center of the colonies (SupFig3a-b). Given that Wnt3a, a direct β-catenin target ^34^, can stabilize GSK3β to prevent pSMAD1 degradation and promote its nuclear accumulation, these findings suggest our wild-type expressing colonies likely over-expressed β-catenin and exhibited enhanced Wnt-dependent differentiation ^35^.

After verifying the functional fidelity of our β-catenin genetically manipulated hESCs, we examined whether preventing tension-dependent phosphorylation impeded mesoderm differentiation. We monitored for mesoderm specification using the nuclear reporter marker, T-mNeonGreen. 24 hours following BMP4 induction, we observed increased numbers of T-positive cells in both control non-transfected hESCs and the hESCs ectopically over-expressing wild-type β-catenin. This contrasted sharply with the lack of mesoderm induction in the hESCs engineered to express the mutant nonphosphorylable β-catenin (mScarlet-NLS+; Fig.1f and g). As such these data indicate that this key domain in β-catenin that is rendered accessible by cellular tension is directly critical for mesoderm specification.

### Y654F mutation prevents Src-dependent β-catenin junctional-nuclear translocation

The kinase Src is activated downstream of cell contractility through phosphorylation by integrin adhesion-associated kinases ^36, 37^. The adhesion-dependent phosphorylated and activated Src in turn phosphorylates β-catenin at Y654 to foster its release from AJs and drive BMP4-dependent mesoderm differentiation ^10, 38^. After confirming that expression of the phosphomutant β-catenin had no impact on tension-induced activation and AJ localization of phospho-Src-family kinases (pSFKs) (Sup.Fig3 a-c), we assessed whether preventing β-catenin Y654 phosphorylation prevented release of β-catenin from the AJs following BMP4 treatment. Consistently, while wild type β-catenin hESCs localized to the regions of high cell contractility showed a significant decrease in AJ localized β-catenin 24 hours following BMP4 stimulation (Fig2 a-c), we did not quantify any discernible decrease in AJ localized β-catenin in the hESCs expressing the phosphodefective mutant β-catenin (see mScarlet-NLS positive cells;Fig2 a-c) ^11^. We also detected higher levels of nuclear-localized β-catenin in the BMP4-stimulated hESCs expressing the wild-type β-catenin as compared to negligible levels detected in the phosphomutant β-catenin expressing cells. These findings confirm that preventing the phosphorylation of β-catenin at the Y654 residue impedes the release of β-catenin from junctional complexes, despite elevated Src family kinase activity, to decrease its subsequent translocation to the nucleus and ultimately impede mesoderm differentiation (Fig2 c-d).

**Fig.2.**
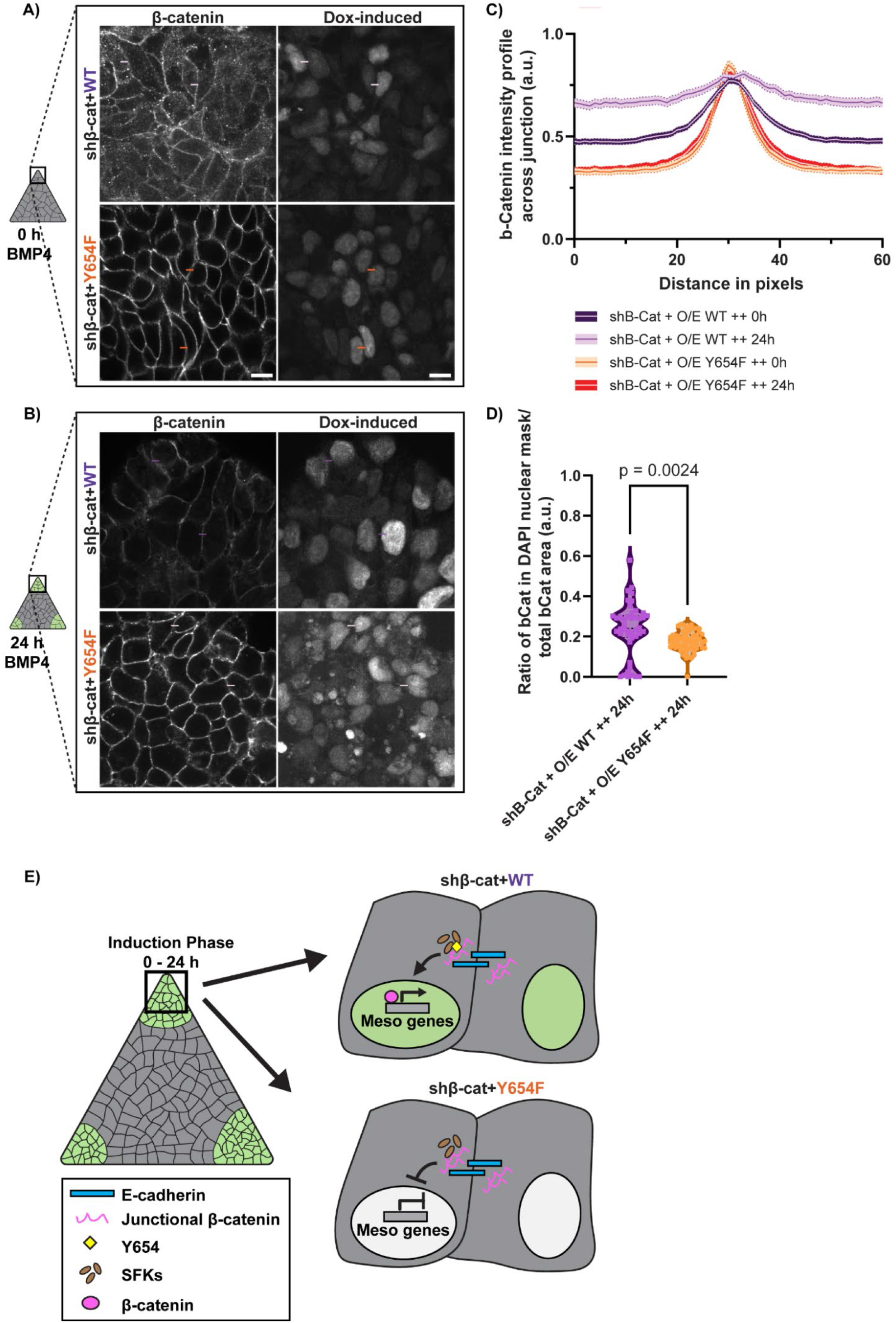
Y654F mutation maintains β-catenin at the junctions and prevent its translocation to the nucleus. A) Representative image of β-catenin staining prior to differentiation in mutant and wild-type cells, with the corresponding expression of the mScarlet-NLS tag of the overexpressing vector. Scale bar = 10 μm. Colored dashes represent examples of areas where the intensity profile was measured across 30 pixels, with the center in the junction, with each color representing the condition shown on the quantification on C. B) Representative image of β-catenin staining after 24h of BMP4 stimulation in mutant and wild-type cells, with the corresponding expression of the mScarlet-NLS tag of the overexpressing vector. Scale bar = 10 μm. C) Quantification of the intensity profiles across junctions for the conditions in A and B. Six profiles were quantified per image for a total of N = (21, 23, 49) for WT-0h, N = (34, 54) for Y654F-0h and N = (7, 15, 22) for WT-24h, N = (65, 41) for Y654F-24h (same samples as in Fig.1D for the 24h timepoint, different channel analyzed). D) Quantification of files shown in B. β-catenin area was thresholded, masked with the mScarlet-NLS StarDist-threshold and normalized to the total mScarlet-NLS area. N = (9, 22, 7) for WT and N= (65, 41) for Y654F cells (same samples as in Fig. 1D, different channel analyzed). Unpaired two-tailed Student’s t-test was performed. E) Schematic of model proposed for mutant x wild-type β-catenin shuttling.

### Y654F mutation prevents epithelial-to-mesenchymal (EMT) transition

We next examined the integrity of β-catenin-dependent signaling targets driving mesoderm expression in the wild-type and mutant β-catenin hESCs ^11, 17^. Consistent with a block in mesoderm specification, 48 hours following BMP4 stimulation, the levels of epithelial-to-mesenchymal transition (EMT) genes including *NCAM1, TWIST* and *SNAI2* ^10, 39^ and the TWIST-induced secreted Wnt inhibitor DKK1^8, 40^ were reduced to varying degrees in the hESC colonies expressing the mutant β-catenin as compared to hESCs expressing the wild-type β- catenin (Fig3b&c). By contrast, levels of the BMP inhibitor *NOG* remained similar in both the wild-type and mutant β-catenin expressing cells following BMP4 stimulation, indicating the BMP pathway remained unaffected by expression of the β-catenin phosphorylation mutation (Fig3d).

**Fig.3.**
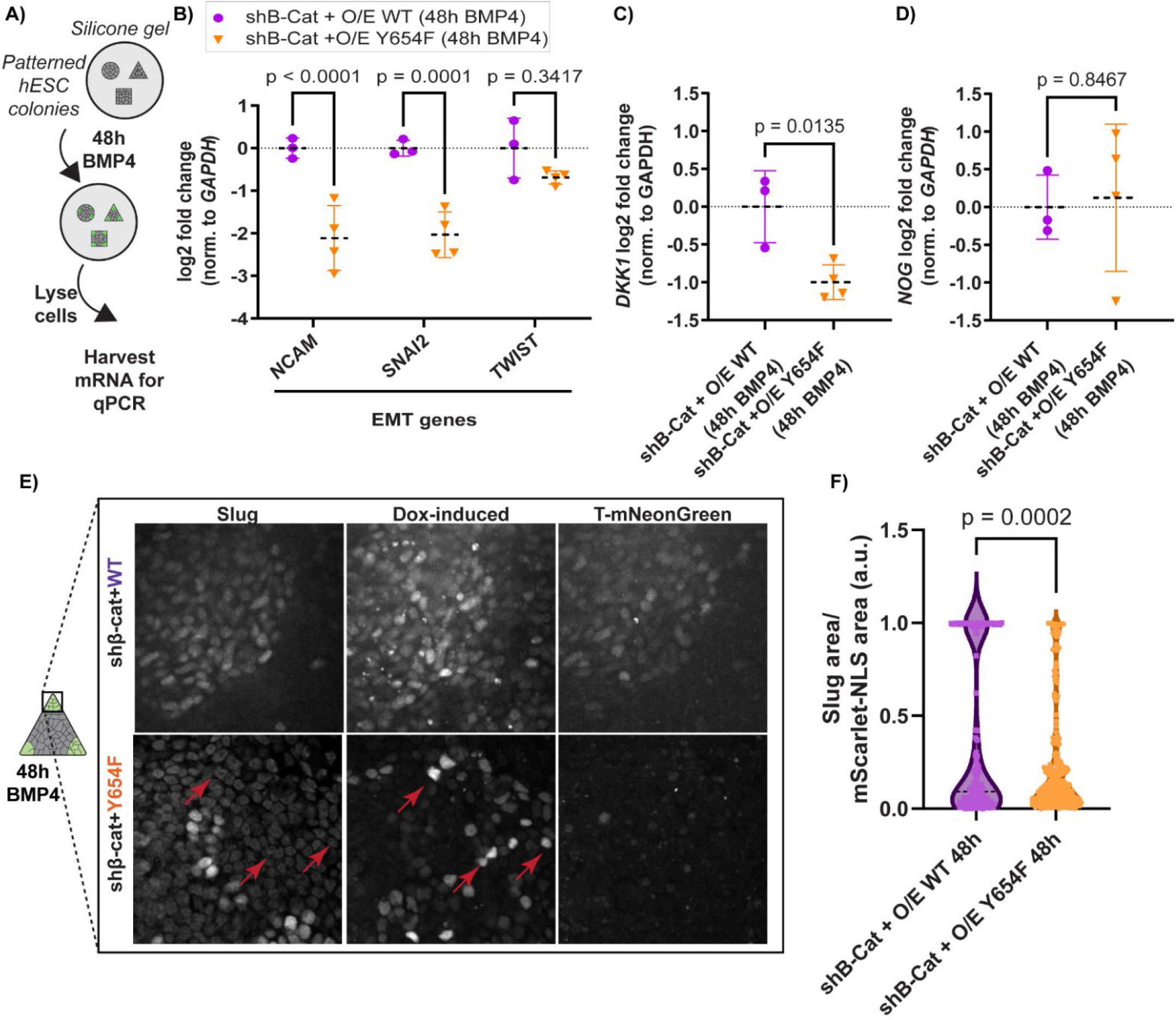
Y654F mutation prevents EMT. A) Schematic of RNA isolation for downstream qPCR application, in which mixed shape wild-type or Y654F mutant colonies were treated with BMP4 for 48h then directly lysed and mRNA was harvested. B) Relative expression levels of EMT genes – *NCAM1*, *SNAI2*, *TWIST*, in wild-type and mutant β-catenin hESCs after 48h BMP4. Data represented as mean (black dashed line) +/- standard deviation (colored full lines) and individual points plotted. N = 3 to 4 independent experiments for each condition. One-way ANOVA test was performed. C) Relative expression of *DKK1* gene, a target of *TWIST*, is higher in wild-type compared to mutant β-catenin hESCs after 48h BMP4. Data represented as mean (black dashed line) +/- standard deviation (colored full lines) and individual points plotted. N = 3 to 4 independent experiments for each condition. Unpaired two-tailed student’s T-test. D) Relative expression of *NOG* gene, a BMP inhibitor, is not changed in wild-type and mutant β-catenin hESCs after 48h BMP4. Data represented as mean (black dashed line) +/- standard deviation (colored full lines) and individual points plotted. N = 3 to 4 independent experiments for each condition. Unpaired two-tailed student’s T-test performed. E) Representative images of mScarlet-NLS β-catenin over-expressing vector expression, T-mNeonGreen and Slug for each condition. Scale bar = 10 μm F) Quantification of images in E). Area of Slug was thresholded using the Triangle auto-threshold and masked with the mScarlet-NLS StarDist-generated threshold to create a mask. Total area was then normalized to the total mScarlet-NLS area. Images were taken from mixed colonies (7C, 6S, 7T) for WT, for a total of N= (52, 52, 53) individual images for WT and (10C, 9S, 8T) for the Y654F mutant, for a total of N= (68, 121, 108) images of the Y654F mutant. Unpaired two-tailed t-test was performed.

Immunofluorescence analysis similarly revealed that the EMT marker Slug (or Snai2) was robustly induced in the wild-type β-catenin expressing hESCs localized adjacent to the T+ cells, indicating that they were undergoing EMT prior to mesoderm specification following BMP4 stimulation ^11^, whereas we detected little to no Slug protein in the hESCs expressing the mutant β-catenin (Fig3e-f). These findings indicate that preventing β-catenin phosphorylation and inhibiting its release from AJs prevents mesoderm specification, as indicated by the failure of these cells to undergo an EMT.

### Mesoderm expression in Y654F mutant can be rescued with Wnt3a exogenous addition or GSK3b inhibition

To ensure the integrity of the Wnt/catenin signaling axis in our mutant β-catenin hESC we conducted a set of mesoderm signaling rescue studies. We treated the hESCs expressing the mutant β-catenin with soluble Wnt3a to determine whether exogenous stimulation of Wnt signaling would promote mesoderm specification in the colonies. Consistent with an intact Wnt signaling axis, addition of exogenous Wnt3a rescued mesoderm differentiation in the mutant β-catenin cells as indicated by induction of T-mNeongreen positive cells (Fig4b) and their concentric spatial distribution within the colonies (Fig4a) ^23^. Interestingly, we also noted that inhibiting GSK3β activity using CHIR99021 (CHIR) similarly rescued T-mNeongreen expression and restored the spatial distribution of mesoderm induction, indicating that preventing β-catenin degradation to artificially elevate cytosolic and nuclear β-catenin can bypass the effects of the mutation on Wnt signaling. We thus arrive at a model in our human gastrulation system where tension-dependent accessibility of AJ β-catenin permits its Src-mediated phosphorylation at residue Y654 β-catenin to facilitate Wnt signaling and direct mesoderm specification is mechanically activated ^11, 19, 20^.

**Fig.4.**
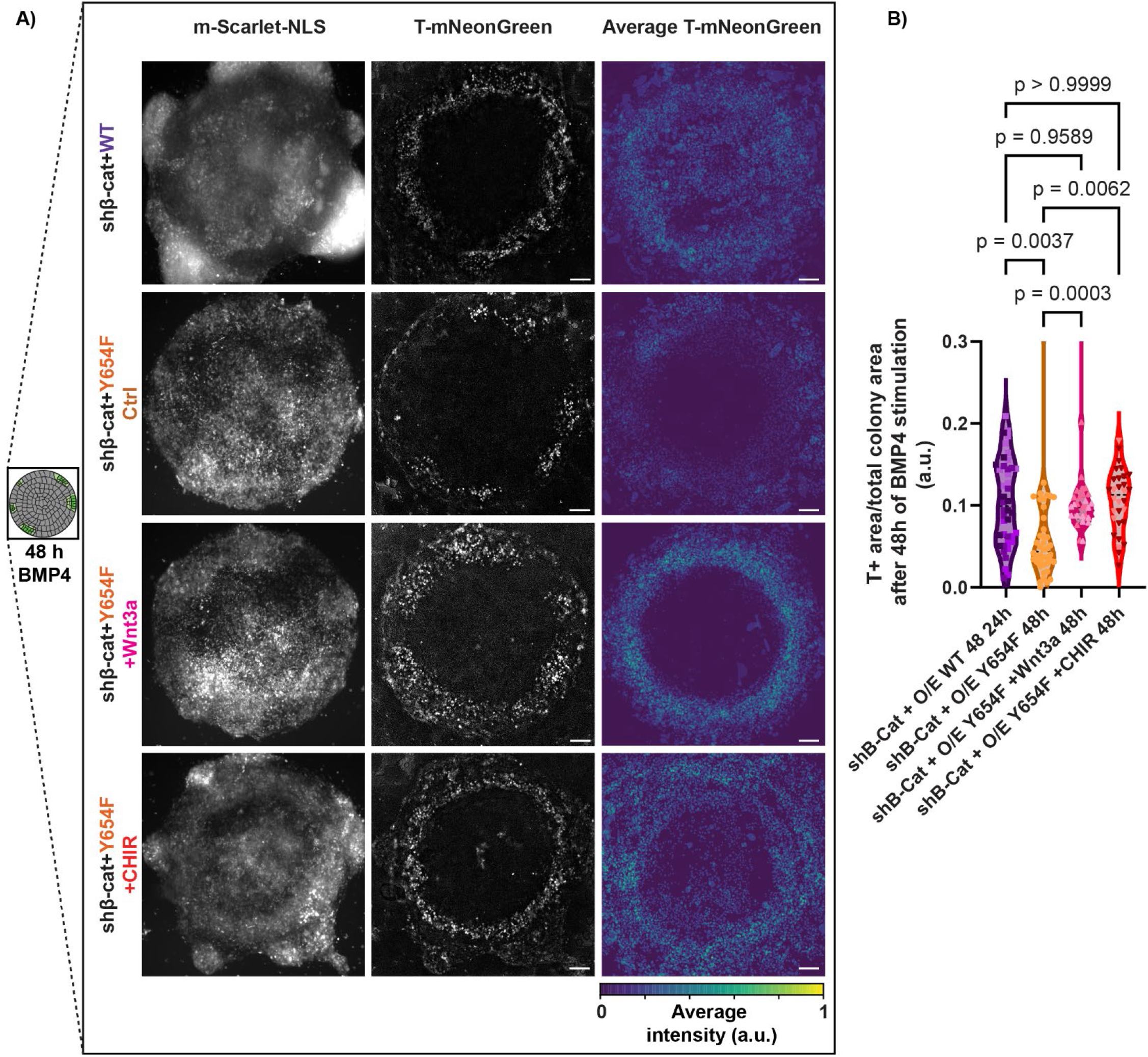
Mesoderm expression in Y654F mutant is rescued by GSK3b inhibition or Wnt3a exogenous addition. A) Representative images of mScarlet-NLS expression (left), T-mNeonGreen (center) and an average of the position of T-mNeonGreen cells in circle patterns for each condition after 48h of differentiation with BMP4. Wnt3a 100 ng/ml and CHIR 2 μm were added throughout differentiation of the mutant. Scale bar = 100 μm B) Quantification of the data shown in A. T-mNeonGreen was thresholded with the StarDist algorithm then masked with the overall shape of the mScarlet-NLS+ area (colony area) and normalized to the total colony area. N = (11, 9, 12,10) for WT, N = (12, 8, 12, 10) for Y654F control, N = (9, 10, 11) for CHIR treatment and N = (12, 12, 12, 5) for Wnt3a treatment. Unpaired two-tailed one-way ANOVA was performed.

## Discussion

Here we provide causal evidence that force-dependent conformational exposure of the Y654 of cadherin-bound β-catenin is a key a mechanism regulating BMP-stimulated Wnt-dependent gastrulation in a hESC gastruloid model. We and others previously implicated mechanical force in EMT induction ^41^ and mesoderm specification during early development ^11^. In prior studies we showed that β-catenin is a mechanosensor that unfolds in response to force and is released through Src-mediated phosphorylation from AJs to be translocated to the nucleus where it functions as a transcription factor to specify Wnt-dependent mesoderm ^19^. Our current results now provide definitive mechanistic evidence linking tissue mechanics and tension to β-catenin Y654 phosphorylation and the EMT and mesoderm specification in this gastruloid model. Our findings are consistent with prior work showing that tension-dependent modulation of β-catenin phosphorylation and Wnt-regulated development is a conserved mesoderm signaling mechanism in a range of species, including zebrafish ^9^, Drosophila ^19^ the cnidarian *N. vectensis* and choanoflagellates ^20^. Homozygous phosphomimetic mutants (Y654E) have been shown to cause embryonic lethality in mice, further implicating the importance of phosphorylation of this residue in higher organism development ^42^. Indeed, mutations close to the Y654 phosphoresidue on β-catenin have been linked to a range of developmental abnormalities ^43^, and introduction of a basic residue close to this phosphodomain causes persistent phosphorylation and sustained Wnt signaling ^44^. Thus, our studies, in combination with prior work, provide concrete evidence for an evolutionarily-conserved mechanically-regulated β-catenin pathway that plays a pivotal role in regulating embryogenesis and tissue development. Beyond the scope of development, our study has major implications for other fields including clarifying the role of tension-dependent phosphorylation of β-catenin in EMT induction during malignancy. Links between tension and Y654 phosphorylation of β-catenin in cancer are indicated by the increased predisposition of heterozygous mice expressing a conditional intestinal-targeted phosphomimetic Y654E mutant to develop intestinal tumorigenesis ^42^, and the mechanical induction of Y654 phosphorylation in a model of mouse colon cancer ^45^.

β-catenin regulates both tissue structure and transcription to modulate development, and is consequently tightly regulated by several post-transcriptional regulatory mechanisms ^13, 14^. β-catenin plays a structural role in adherens junctions through its association with E-cadherin and, in the nucleus, drives transcription through its binding to TCF/LEF. Conversely, cytosolic pools of β-catenin bind to the AXIN/APC destruction complex which directs its degradation ^46^. These diverse interaction partners have overlapping domains that competitively bind to the β-catenin central groove, which mediates context-dependent spatial segregation ^47^. Not surprising, these functional interactions are refined by either N-terminal or C-terminal phosphorylation ^46^. Of note, sequential N-terminal domain phosphorylation of the cytosolic β-catenin/APC/AXIN occurs first by CK1α at S45 then by GSKβ at T41, S37 and S33, targeting β-catenin for degradation ^48^. In addition to these modifications, there exist feedback mechanisms to control β-catenin function, as indicated by the inhibition of GSK3β-dependent degradation of β-catenin that stabilizes its cytosolic levels and increases the pool of activated nuclear β-catenin through secreted Wnts^49, 50^. Interestingly, molecular dynamic simulations and experimental evidence indicate that junctional tensile forces provide the needed context to expose the C-terminal helix domain where phosphorylation of Y654 occurs, toggling junctional β-catenin to facilitate its nuclear translocation and transcriptional activity^19^. Whether these two modes of regulation represent distinct pools of β-catenin or if they directly interact is currently under investigation ^48, 51^. Our studies showed that while we retained junctional β-catenin pools and repressed nuclear transcriptional activity through expression of the Y654F phosphomutant, we were able to rescue Wnt-dependent mesoderm signaling by addition of Wnt3a and by inhibiting GSK3β-dependent β-catenin degradation with CHIR99021. This result supports the concept that there are separately regulated pools of cytosolic and adhesion-associated β-catenin. Nevertheless, given that we used a dual knock-down of endogenous β-catenin combined with its over-expression, our studies could not discern whether the GSK3β inhibition rescue was mediated by increased availability of endogenous cytosolic β-catenin that was not fully depleted through our knockdown strategy, or was mediated via a pool of redirected cytosolic mutant β-catenin that independently stimulated mesoderm signaling. Although this possibility would need to be explored through the use of an endogenous Y654F β-catenin mutant created through manipulation with a CRISPR point mutation ^52^, our studies do concur with the concept that there exists independently regulated pools of AJ-associated and cytosolic β-catenin.

Morphogens and growth factors play important roles during embryogenesis and tissue development. Central to gastrulation is the role of BMPs in driving endogenous Wnt and NODAL secretion to govern cell lineage specification ^53, 54^. Nevertheless, early embryogenesis and tissue development are also characterized by force-generating tissue and cell movements. Mechanistically, morphogenesis is known to occur through the activation of Wnt/PCP ^3^ and Rho1/MyoII^2^, triggering invagination of the mesendoderm layer and primitive streak formation during gastrulation^4, 5, 55^. Part of the complexity of gastrulation arises from morphogenesis and pattern formation occurring simultaneously in the embryo. While it is known that cell fate determination can induce shape changes and migratory behavior^56^, the converse has only recently been demonstrated^9–12^. Our data reveal how tissue mechanics regulates development by mechanically-activating β-catenin to permit its functional phosphorylation that is necessary for driving Wnt signaling. These findings reveal how morphogenetic forces not only sculpt tissue organization but also modify morphogen signaling to regulate cell fate, providing a new and as of yet under-appreciated perspective on how contextual cues synergistically modify diffusible morphogen signals.

Recently established three-dimensional (3D) organoid models that recapitulate various aspects of cell dynamics during development including human post-implantation embryogenesis ^57, 58^, neural tube and trunk formation ^59–61^ and forebrain development ^62^ have underscored the importance of synergism between spatial cues and morphogens in tissue-specificity ^63^ . Our hESC gastruloid model, is a model that not only recapitulates key features of early embryogenesis but also provides a tractable system to test the interplay of mechanical force regulation and spatial cues in morphogen signaling ^11, 63, 64^. Our model builds upon and extends prior work in *in vitro* gastruloids, and exploits the use of micropatterned gastruloids to address the role of mechanical forces and spatial patterning, as was reported for neuroectodermal tissue ^65^. The versatility of our gastruloid model and the relative ease with which it can be manipulated lends itself to future studies aimed at exploring the role of mechanical forces in other germ layer specification programs, including addressing synergism with the Activin/Nodal pathway and impact on endodermal and extraembryonic lineages.

## Acknowledgments

This work was supported by NIH NCI grants R35CA242447 (V.M.W.) and R01CA222508 (V.M.W.). N.M.E.A was additionally supported by the Schlumberger Foundation – Faculty for the Future Fellowship, the UCSF Michael Litchman Discovery Fellowship, and the UCSF Lloyd Kozloff Fellowship. We acknowledge the Biological Imaging Development Center (BIDC) at UCSF and in particular the use of the SoRa microscope system, supported by the NIH shared equipment grant S10OD028611-01. We also acknowledge the PFCC (RRID:*SCR_018206*) supported in part by Grant NIH P30 DK063720 and by the NIH S10 Instrumentation Grant S10 1S10OD021822-0. We also thank Jeffrey Bush and Ophir Klein for their valuable comments on the manuscript.

## Author contributions

Conceptualization, N.M.E.A, J.N.L and V.M.W; Methodology, N.M.E.A and J.N.L.; Investigation, N.M.E.A.; Writing – Original Draft, N.M.E.A..; Writing –Review & Editing, N.M.E.A., J.N.L., V.M.W..; Funding Acquisition, N.M.E.A., V.M.W.; Resources, N.M.E.A., J.N.L., A.G., A.J.E., V.M.W.; Supervision, J.N.L, V.M.W.

## Declaration of interests

The authors declare no competing interests.

## Supplemental information titles and legends

**Sup.Fig1.**
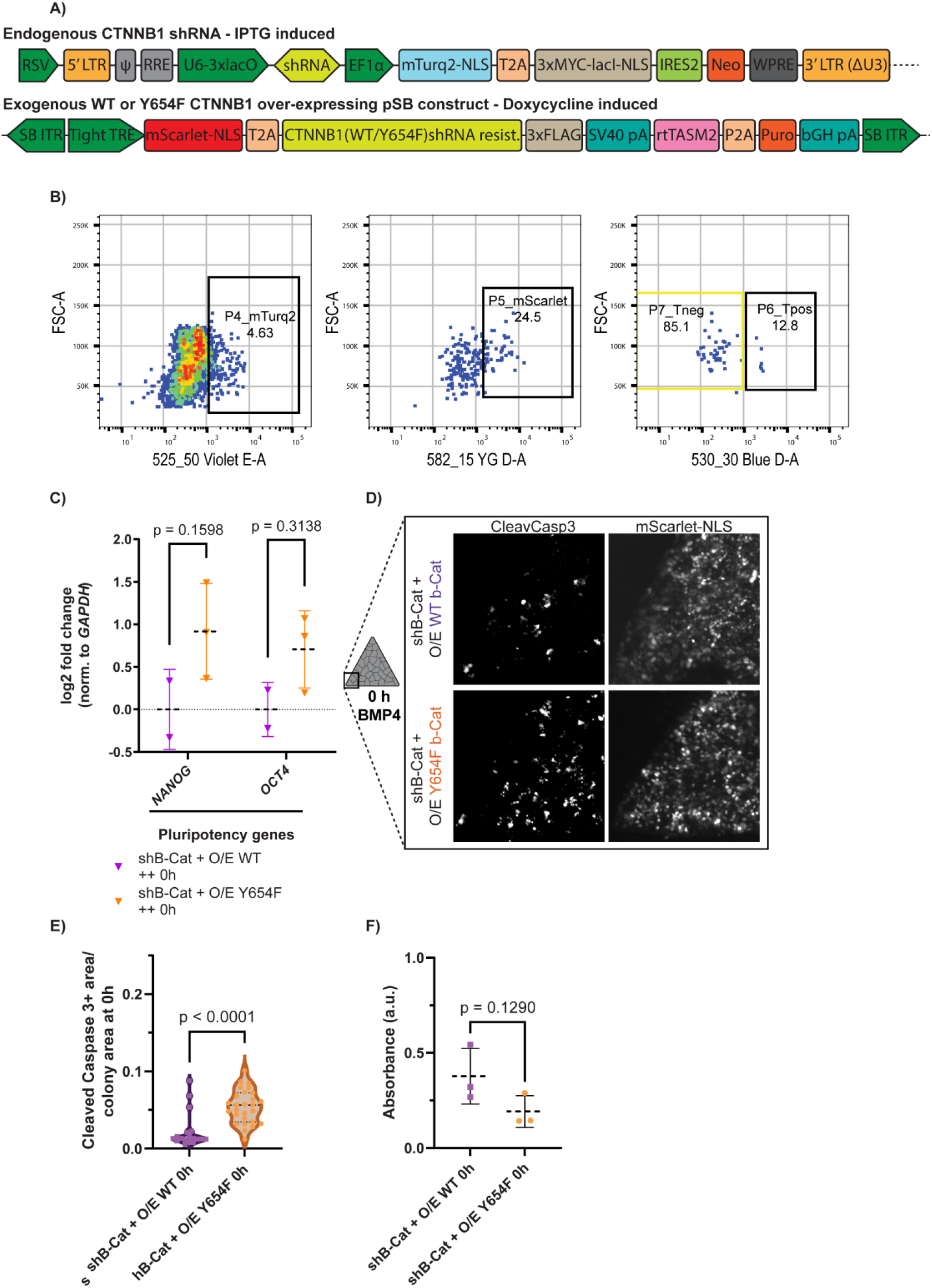
Mutant cell line Y654F β-catenin was created with two integrated constructs and maintains pluripotency and proliferation. A) Features of the constructs used for this study. IPTG-Inducible shRNA targeting *CTNNB1* (gene encoding for β-catenin) contains an mTurq2 tag with a nuclear localization signal (NLS) and three MYC tags constitutively expressed in cells infected with this construct. Doxycycline (DOX)-inducible over-expressing Sleeping Beauty transposon vector contains the sequence for either the wild-type *CTNNB1* or Y654F mutated *CTNNB1*, flanked by an mScarlet tag with a nuclear localization signal on the 5’ end and three Flag tags on the 3’ end. B) Representative images showing fluorescent activated cell sorting (FACS) gating strategy. After size gates, selected cells were sorted for high knockdown expression (P4_mTurq2-NLS), medium high overexpressing vector (P5_mScarlet-NLS) and negative expression of T-mNeonGreen (P7_Tneg). C) Relative expression levels of pluripotency genes – *Nanog, OCT3/4 –* in wild-type and mutant β-catenin hESCs prior to BMP4 stimulation. Data represented as mean (black dashed line) +/- standard deviation (colored full lines) and individual points plotted. N = 2 (WT) or 3 (Y654F) independent experiments for each condition D) Crystal Violet absorbance assay to assess proliferation of wild-type and mutant cell lines (N=3 for both). Unpaired two-tailed Student’s t-test was performed on data. E) Representative grayscale image of Cleaved-Caspase-3 staining (left) and m-Scarlet-NLS (right) expression in wild-type and mutant colonies before the start of differentiation (0h). Scale bar = 100 μm F) Quantification of the data displayed in E. The background of Cleaved-Caspase-3 images was subtracted, then the images were thresholded and masked with the colony area and normalized to the total colony area. N = (5, 6, 6) for WT, N = (7, 7, 8) for Y654F control. Unpaired two-tailed Student’s t-test was performed.

**Supp.Fig2.**
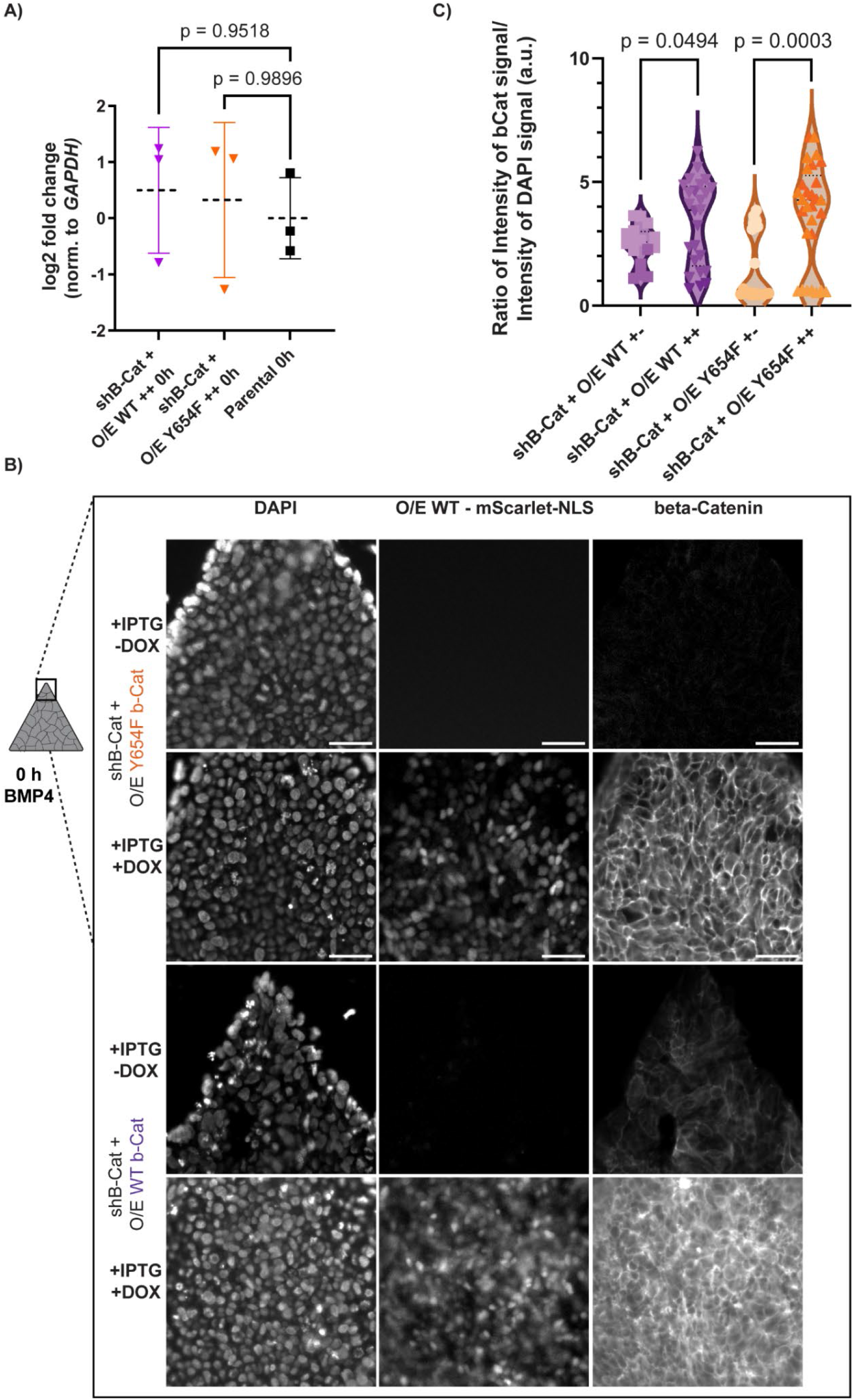
β-catenin is normally expressed at adherens junctions in wild-type and mutant cell lines. A) Relative expression levels of *CTNNB1* in parental, wild-type and mutant β-catenin hESCs prior to BMP4 stimulation. Data represented as mean (black dashed line) +/- standard deviation (colored full lines) and individual points plotted. N = 3 (Parental), 2 (WT) or 3 (Y654F) independent experiments for each condition B) Representative images of colonies of hESCs plated according to Fig.1A prior to differentiation with and without DOX overnight induction. mScarlet-NLS on the center indicates cells expressing the DOX-induced construct, while right panels show β-catenin resulting expression, with the left panel indicating DAPI nuclear signal. Scale bar = 50 μm C) Quantification of the data in B, where loss of β-catenin intensity with the knockdown only (+IPTG-DOX) was quantified against the knockdown + overexpression strategy for wild-type and mutant. Each datapoint represent the intensity of the β-catenin signal in a colony normalized to the overall DAPI signal. Number of images is N = (1,11,5) for WT +-, N = (11,7) for Y654F +-, N = (3,10,12) for WT ++, N = 12,10,7) for Y654F ++.

**Supp.Fig3.**
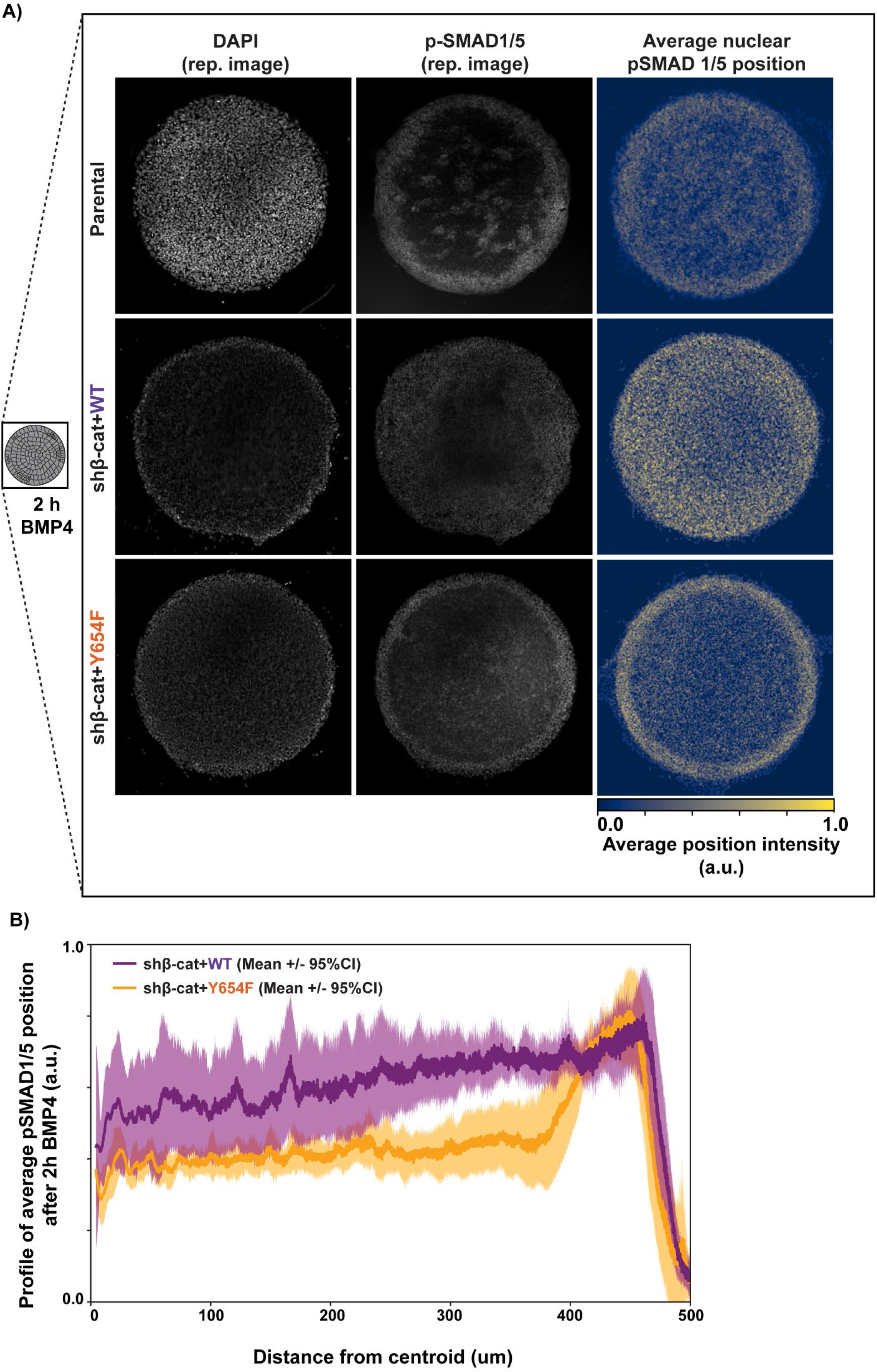
pSMAD1/5 nuclear expression distribution after 2h of BMP4 differentiation is not affected by this mutation. A) Representative grayscale image of mScarlet-NLS (left) and p-SMAD1/5 (center) expression in parental, wild-type and mutant colonies after 2h of BMP4 stimulation. Average nuclear p-SMAD1/5 position image (right) was created averaging pSMAD1/5 images after threshold with the StarDist algorithm. N = (8,7,7,3) circle parental colonies, N = (4,3,4) circle colonies for WT images and N= (4,4,3) circle colonies averaged for mutant colonies. B) Profile quantification of pSMAD1/5 threshold. Pixel intensity of each pSMAD1/5 threshold image shown above was measured from the centroid to its edge. Plot displays the average with a mean filter of 20 pixels (dark line) and 95% CI of N = (4,3,4) for WT and N= (4,4,3) for mutant.

**Supp.Fig4.**
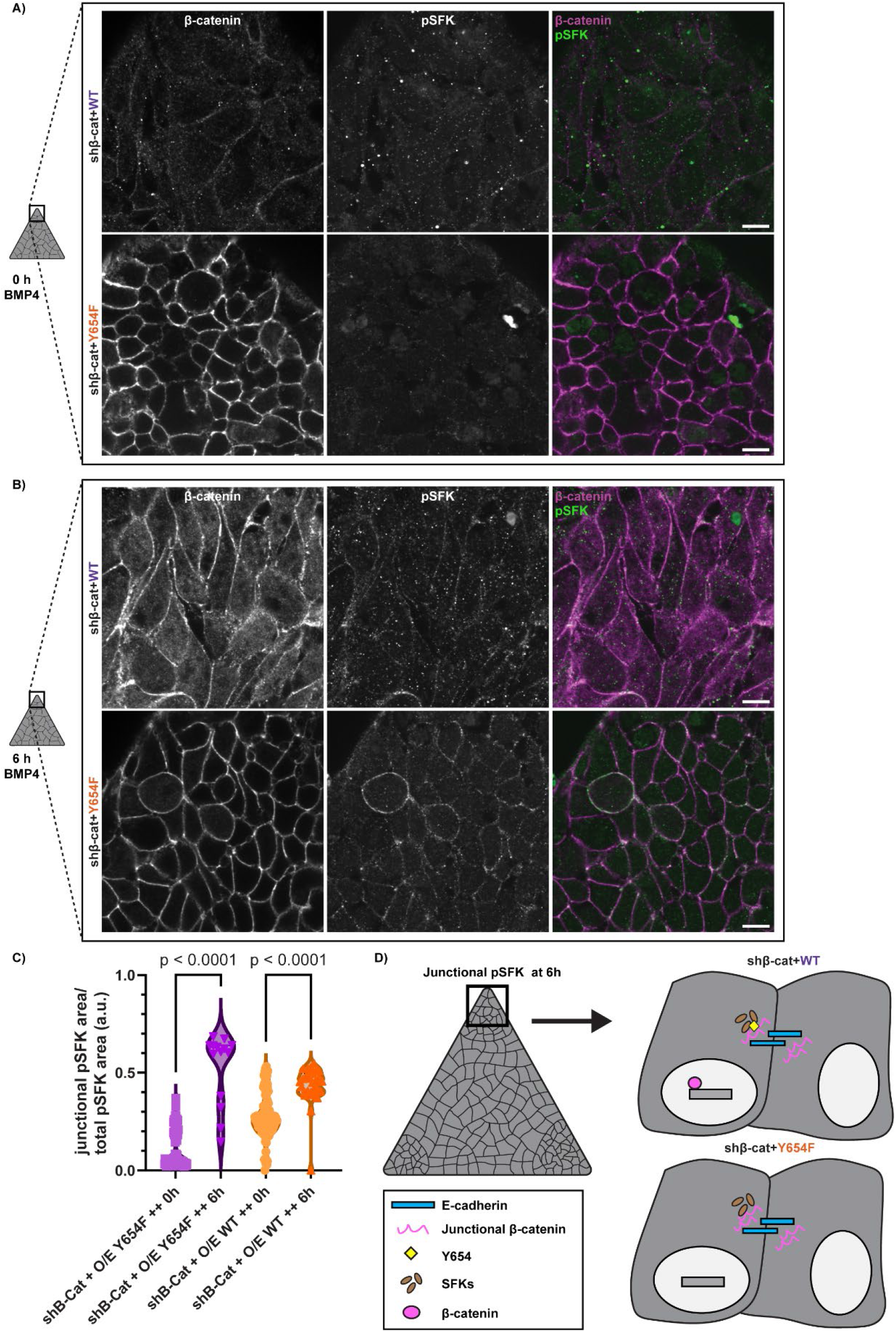
pSFK recruitment to β-catenin junctions during differentiation is not affected by this mutation. A) Representative images of β-catenin (left), pSFK (center) staining, and the merged image of both (right) prior to the start of BMP4 differentiation. B) Representative images of β-catenin (left), pSFK (center) staining, and the merged image of both (right) after 6h of BMP4 stimulation. C) Quantification of data in A and B. Area of pSFK was thresholded using the Triangle auto-threshold and masked with a β-catenin thresholded image. Masked pSFK was then normalized to the total pSFK thresholded area. Images were taken from the corners of mixed colonies for a total of N= (15, 32, 17) individual images for WT at 0h, N= (16, 16, 35) for WT after 6h of BMP4 stimulation, N= (19, 33) images of the Y654F mutant at 0h, N = (24, 31) for the Y654F mutant after 6h of BMP4 stimulation. D) Cartoon indicating that at 6h, SFKs are active and localized to the junctions in the corner of these colonies, but only in WT cells can β-catenin be phosphorylated at the Y654 residue, removed from the junction, and translocate to the nucleus.

## Tables with titles and legend

**Table S1.**
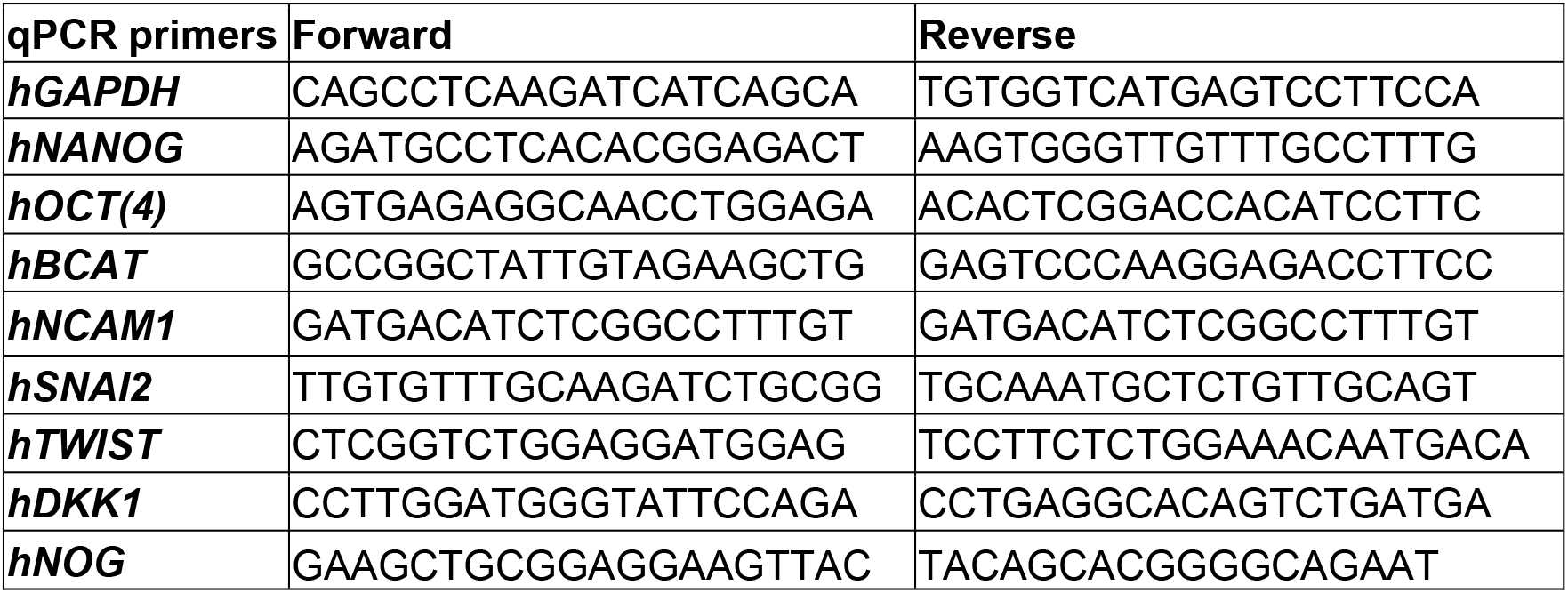
List of primers used in this study.

## STAR Methods

### Key Resources Table

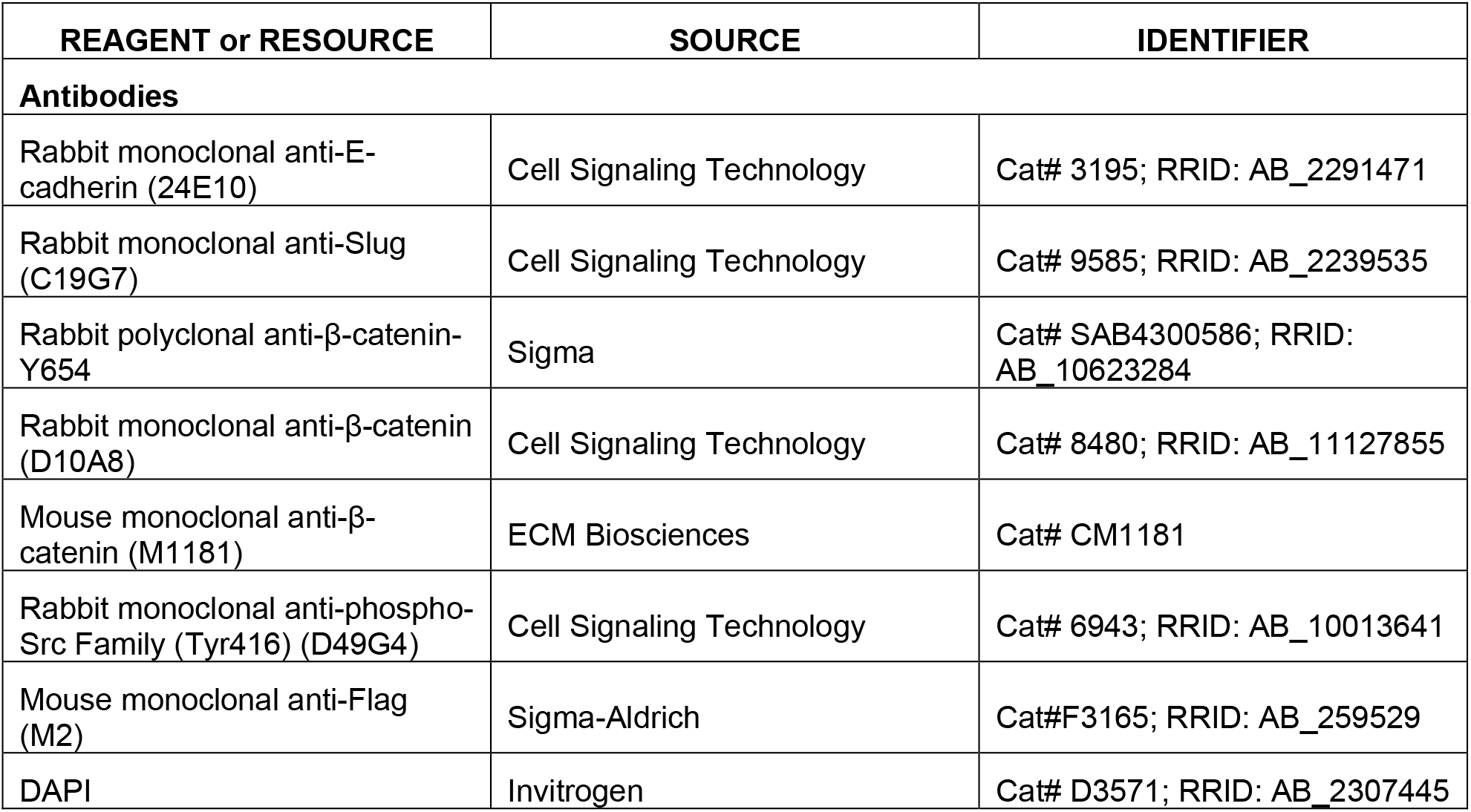
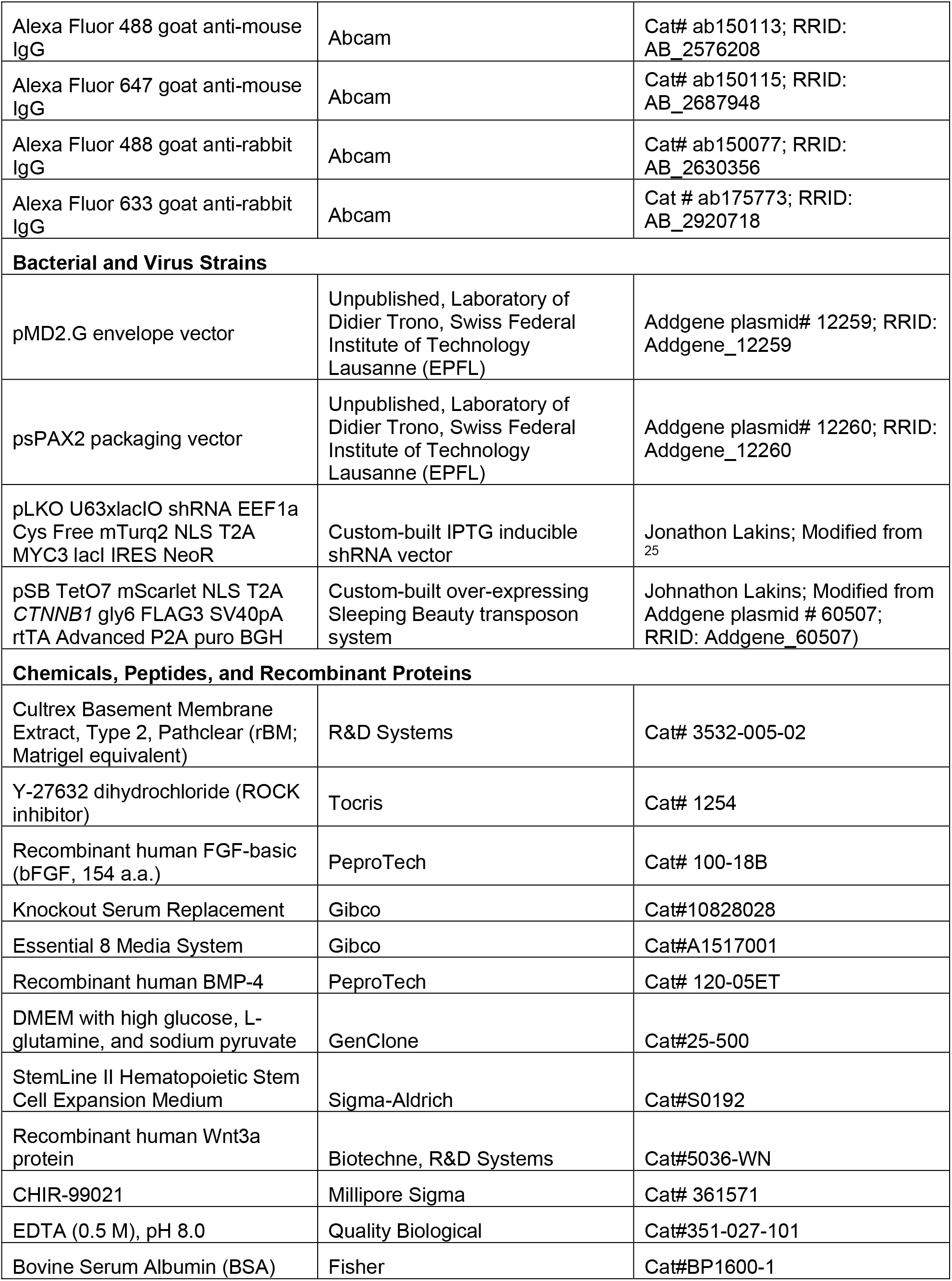
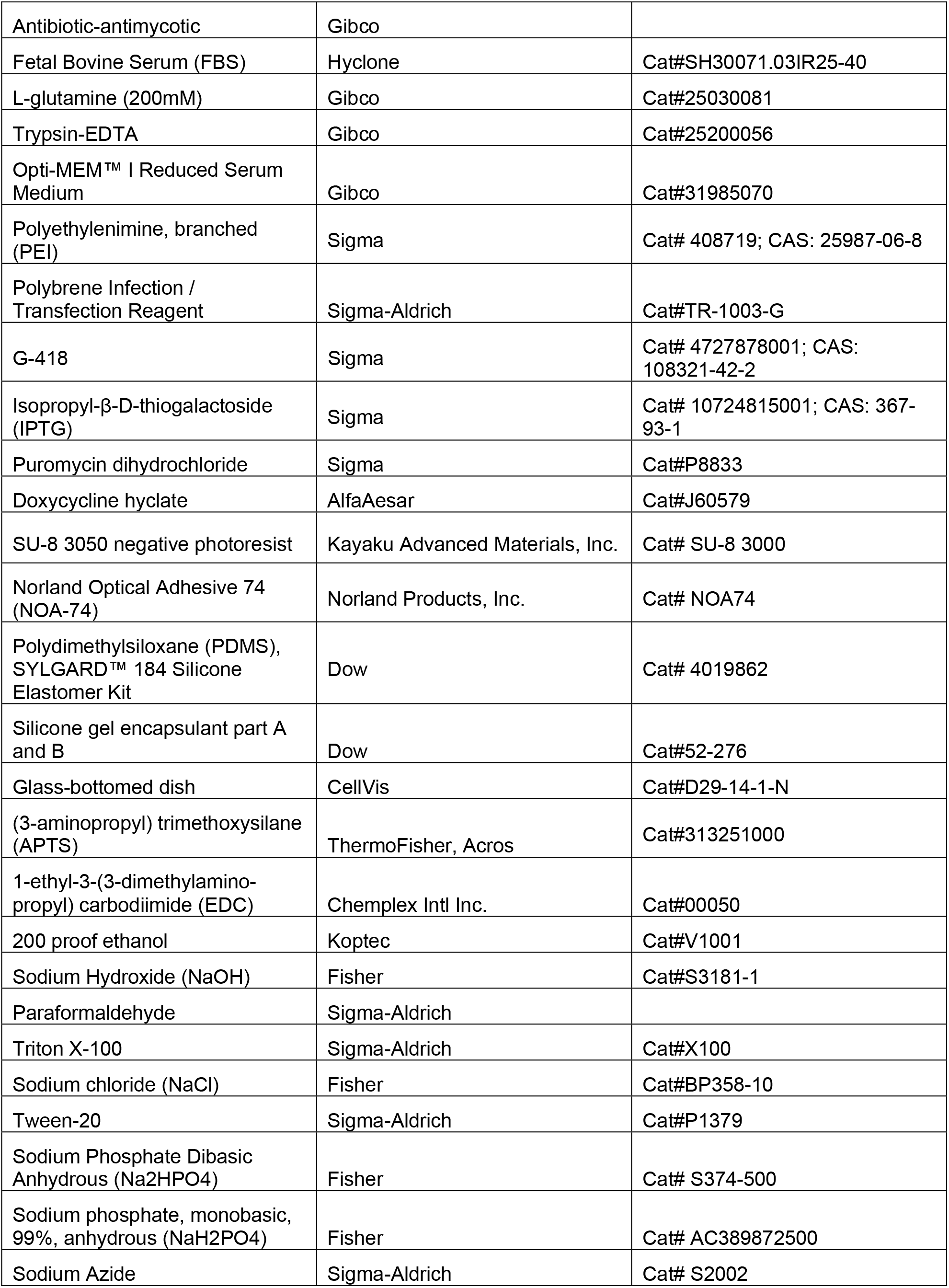
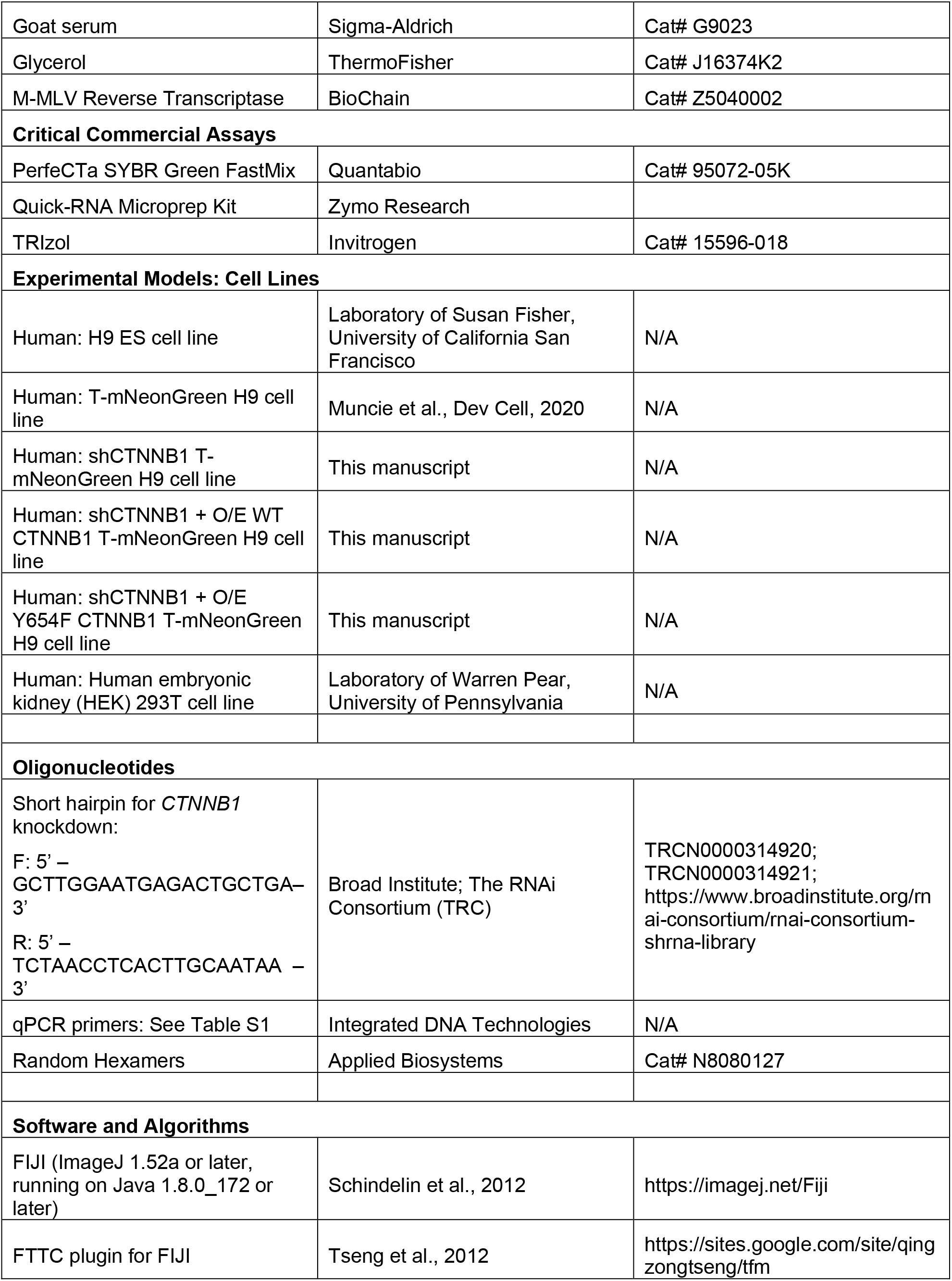
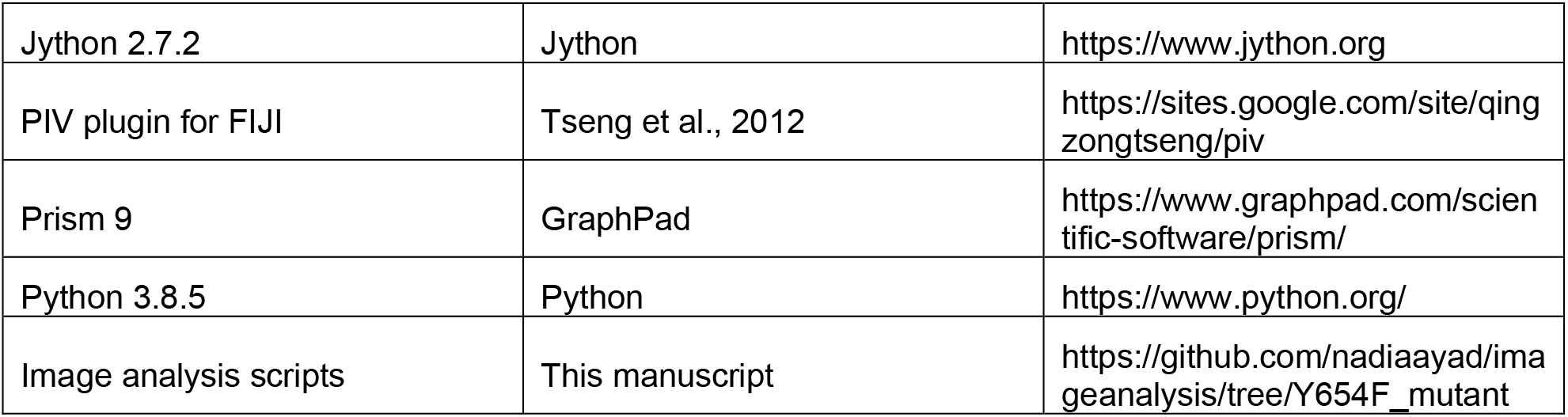

### Resource Availability

#### Lead Contact

Further information and requests for resources and reagents should be directed to and will be fulfilled by the Lead Contact, Valerie M. Weaver.

#### Materials Availability

Modified plasmids are deposited on AddGene.

#### Data and Code Availability

The code generated during this study for data analysis is publicly available on GitHub: https://github.com/nadiaayad/imageanalysis/tree/Y654F_mutant.

### Experimental Models and Subject Details

#### Cell Lines

The parental cell lines for this study are human embryonic stem cells (H9, female) obtained as a gift from the Laboratory of Susan Fisher at UCSF. T-mNeonGreen reporter cell lines were generated as described in ^11^.

All cell lines were regularly maintained in feeder-free conditions on tissue culture plastic coated with reconstituted basement membrane extract (rBM; Matrigel equivalent; R&D systems) DMEM with high glucose, L-glutamine, and sodium pyruvate (GenClone). They were cultured in a 50% complete Essential 8 (E8) media (Gibco) and 50% KnockOut^TM^ Serum Replacement (KSR) media mix (Gibco) with 10 ng/ml bFGF (PeproTech) and 1x antibiotic-antimycotic (Gibco), referred to as hESCs growth media. Media was replaced every 24h and cells were passaged every 3-4 days with 0.5 mM EDTA/2% BSA (Fisher) in PBS with 50% E8/50% KSR media supplemented with 10 μM Y-27632 (ROCK inhibitor; TOCRIS). After 24h post-passaging, the cells were replaced with 50% E8/50% KSR media without Y-27632.

Human embryonic kidney (HEK) 293T cell lines were obtained as a gift from the Laboratory of Warren Pear at University of Pennsylvania and used for transfection and production of lentivirus. Maintenance media for HEK 293T cells was DMEM with high glucose, L-glutamine and sodium pyruvate and supplemented with 10% fetal bovine serum (FBS; Hyclone), 4 mM L-glutamine (Gibco) and 1x antibiotic-antimycotic. Upon reaching 75% confluency, cells were passaged with 0.05% trypsin-EDTA (Gibco).

All cell lines were maintained in a humidified incubator at 37°C°/5% CO_2_ and evaluated routinely for mycoplasma contamination and confirmed to be negative.

Experiments involving hESCs were approved by the University of California San Francisco Human Gamete, Embryo and Stem Cell Research Committee (UCSF GESCR) in the study numbered 11-05439.

#### Generation of β-catenin knockdown cell line

A validated shRNA (TRCN0000003845 GCTTGGAATGAGACTGCTGAT, The RNA Consortium; TRC) targeting human β-catenin was cloned into a custom modified version of the TRC U6 inducible 3 x LacO vector pLI-913. The human EEF1a promoter replaced the hPGK promoter in pLI-913 to drive expression of an IRES mediated polycistonic mRNA consisting of an upstream polyprotein of mTurq2 (SV40) NLS and myc3 tagged lacI repressor (SV40) NLS separated by a T2A peptide and a downstream Neomycin phosphotransferase for selection with G418 (Supl.Fig 1). Prior to transfection, HEK 293T cells were plated at a density of 0.8 × 106 cells in a 35 mm tissue culture plastic dish. Twenty-four hours later HEK 293T cells were washed gently with PBS to remove serum and media was replaced with Opti-MEM media (Gibco). For transfection, 0.33 μg of psPAX2 packaging vector (Addgene) with 0.16 μg pMD2.G envelope vector (Addgene) and 0.55 μg shRNA vector, for a total of 1 μg of DNA, were mixed with Opti-MEM media in a 100 μl volume for a 5-minute incubation at room temperature. Parallelly, 3 μg of polyethylenimine (PEI; Sigma) was mixed in Opti-MEM media to a total volume of 100 μl and incubated for 5-minutes at room temperature. Then, both solutions were combined in a microcentrifuge tube and incubated or an additional 15-minutes at room temperature before being mixed into the HEK 293T cells in the 35 mm dish incubated in Opti-MEM media. After 6h post-transfection, media was replaced for normal HEK 293T growth media supplemented with 200 μM IPTG.

Forty-eight hours post-transfection, viral supernatant was collected and centrifuged 2x at 500x g, carefully collecting the supernatant and discarding the pellets both times to remove remaining detached 293T cells. Centrifuged supernatant was then supplemented with 4 μg/ml polybrene (Sigma). T-mNeonGreen reporter cell lines cultured on rBM-coated tissue culture plastic were immediately transduced with a 1:4 mix of the supernatant: hESCs growth media for 24h. Media was then replaced with fresh hESCs growth media for an additional 24h, prior to selection with 200 μg/ml G-418 (Sigma). During maintenance culture of these cells, 200 μg/ml of G-418 (Sigma) was continuously added to the hESCs growth media to prevent loss of inducible *CTNNB1* knockdown. Knockdown of *CTNNB1* was induced with 0.2 mM isopropyl-β-D-thiogalactoside (IPTG; Sigma) during seeding on gels.

#### Generation of mutant and wild-type cell lines

The complete wild type human β-catenin coding sequence was obtained from the DNA plasmid repository of Arizona State University (DNASU) (Clone HsCD00079878). Golden Gate Assembly of PCR amplified fragments using the Type IIS restriction enzyme Esp3i was used to simultaneously prepare silent mutations of the shRNA recognition sequence, introduce the Y654F mutation and prepare a polyprotein with an upstream mScarlet (SV40) NLS separated from β-catenin using a T2A peptide. Wild-type and Y654 mutants were cloned into sfiI digested pSBtet-Pur (Addgene plasmid # 60507; RRID:Addgene_60507) ^26^ adding a C-terminal 3xFlag epitope tag and replacing the firefly luciferase gene (Supl.Fig 1).

Prior to transfection, *CTNNB1* knockdown T reporter H9 were trypsinized from rBM-coated tissue culture plastic, resuspended in DMEM with 10% FBS and recovered by centrifugation. Pelleted cells were washed once in 0.5 ml of Electroporation Buffer (20 mM HEPES-10.5 mM NaOH, 125 mM KCl, 2 mM MgCl_2_, 0.5% (w/v) Ficoll 400 pH 7.6) and recovered by centrifugation in a 1.5 ml eppendorf. Pelleted cells (∼20 µl) were resuspended in ∼0.16 ml Electroporation Buffer and combined with 10 µl of a 20X mix of ATP and reduced glutathione (20X stock is 40 mM ATP, 100 mM reduced glutathione in 1x Electroporation Buffer; pH adjusted to 7.6), 2.5 μg of the transposase vector SB100x ^66^ (Addgene plasmid #34879; RRID: Addgene_34879) and 5 μg of *CTNNB1* WT or Y654F Sleeping Beauty vector in 10 µl of ddH_2_O. The cell DNA suspension was transferred to a 0.4 cm electroporation cuvette (Biorad) and electroporated by exponential decay on a Biorad Gene Pulser II using a capacitance of 500 µF and total voltage of 200 mV. After 5 minutes of post transfection incubation at room temperature, transfected cells were recovered by resuspension in 1 ml DMEM with 10% FBS and centrifugation. Pelleted cells were resuspended in hESC growth media supplemented with 10 ng/ml bFGF (PeproTech) and 10 μM Y27632 (Tocris) and replated on rBM coated 35mm plates. After 24 hours, media was replaced for an additional 24 hours, prior to selection with 1 μg/ml puromycin for 4 days (Sigma). During maintenance culture of these cells, 200 μg/ml of G-418 (Sigma) and 1 μg/ml puromycin (Sigma) was continuously added to the hESCs growth media to prevent loss of inducible *CTNNB1* knockdown and overexpressed *CTNNB1.* A pure population of cells expressing both the knockdown and overexpressed *CTNNB1* was obtained by transiently inducing the overexpressed vector with 1 μg/ml doxycycline (AlfaAesar) for 3 hours and subsequently using Flow Activated Cell Sorting (FACS) to purify it the next day, by gating for high mTurq2 (knockdown vector) and medium-high mScarlet (overexpressing vector) expression, as described in the main text. Single cells isolated by FACS were plated on rBM-coated tissue culture plastic in hESC media 10 μM Y27632 (Tocris). Fresh hESC media with Y27632 and 200 μg/ml of G-418 (Sigma) and 1 μg/ml puromycin (Sigma) was added every 24 hours until colonies of undifferentiated hESCs were visualized.

#### Fabrication of Silicone Gels

In order to use a soft substrate that simulate physiological stiffness and also could be used in high resolution microscopy, we used silicone (e.g. polydimethylsiloxane) gels of tunable stiffness as described in ^29^ as a substrate in our studies. Briefly, we used 29 mm glass bottom dishes (CellVis) designed for high resolution imaging and cleaned them with a 3N NaOH solution (Fisher) for 20 minutes. Dishes were thoroughly washed 5x with distilled water, then dried under a N_2_ (Airjet) air jet, to avoid any debris on the surface. We mixed the silicone gel (DowSil) at a 1:1 polymer A: crosslinker B ratio so that the final shear storage modulus would be at 1.5 kPa. The uncured gel was then degassed in a vacuum chamber. Following degassing, the 29 mm glass-bottomed dried dish was spin-coated (VTC-100 from MTI Corporation with an AP-1400C/V AutoBo Elec. Technology Limited vacuum pump) at a 10sec/1000rpm and 60 sec/7500 rpm program with 100 μl of the gel on its center and then baked at 80°C for 2h.

Silicone gels for traction force microscopy were generated similarly but with the addition of custom-synthesized PMMA far-red beads as described in ^67, 68^, in a 1:100 ratio in the uncured mixed gel.

#### Patterning of Silicone Gels

Stencils were generated as described in ^30^. Briefly, polydimethylsiloxane (PDMS; Dow) stamps with designed geometric patterns were fabricated on custom silicon wafers generated using negative photoresist (SU-8; Kayaku Advanced Materials Inc.). Stencils were created by placing the PDMS stamps on a flat slab of PDMS and then wicking a UV-curable adhesive NOA-74 between the stamp and slab, forming a stamp/glue/slab “sandwich”. In order to cure the adhesive, the ”sandwich” was placed on a medium wavelength UV source (Spectroline EN-180, 306 nm peak) for 5 min with the stamp on the top position, then the “sandwich” was turned, so that the slab could be in the top position, for an additional 5 min UV-exposure, to ensure full curing of the adhesive.

Next, stencils were placed on the cured silicone gel substrates gently as to not disturb the surface of the gel but ensuring that there are no gaps between the stencil and gel. To conjugate ECM proteins to the surface in a pattern, gels with stencils were incubated with 95% ethanol (Koptec) and 0.5% (3-aminopropyl) trimethoxysilane (APTS; Sigma-Aldrich) for 10 min. At this point, dishes were taken to a sterile cell-culture hood, then sprayed with 70% ethanol. Solution was then aspirated, and gels were washed with PBS 3x. A solution with 100 μg/l of 1-ethyl-3-(3-dimethylamino-propyl) carbodiimide (EDC, TCI America) was mixed with rBM in PBS and placed on the gels and left at 4°C/overnight or at 37°C/1h.

Following the conjugation of ECM, gels were then washed with PBS 3x to remove remaining unconjugated rBM. Using sterile, autoclaved forceps, stencils were gently removed, making sure that surrounding gel was not disturbed. At this point, gels could be stored for 1 week at 4°C or immediately used for cell seeding.

#### Plating hESCs on patterned silicone gels

Cells were passaged by washing the hESCs cultured on rBM-coated plastic 3x with DMEM, then incubating the cells with 0.05% trypsin-EDTA supplemented with 10 μM Y-27632 for 5 min at 37°C. Trypsin was quenched by adding the same volume of hESCs growth media supplemented with 10 μM Y-27632 to the dish. Cells were centrifuged and resuspended in hESCs growth media supplemented with 10 μM Y-27632 and counted using a hemocytometer. Patterned hESCs colonies were obtained by placing 150,000 cells on the patterned silicone gels, making sure that the cells only cover the glass bottomed well (about 400 μl), without spillage to the plastic part. For the sh*CTNNB1* +/- WT or Y654F overexpression vector cell line experiments, 0.2 mM IPTG was added to the media upon seeding and maintained throughout the course of the experiment. The next day the media was changed to hESCs growth media without Y-27632. The following day, the media was changed to hESCs growth media only and with 1 μg/ml doxycycline (DOX; AlfaAesar) overnight for experiments with the induction of the over-expressing vector WT or Y654F β-catenin.

#### BMP4 Differentiation

Differentiation was induced by replacing the hESCs growth media with StemLine II Hematopoietic Stem Cell Expansion Medium (Sigma) supplemented with 10 ng/ml bFGF (PeproTech) and 50 ng/ml BMP4 (PeproTech), thereafter called differentiation media. For cell lines with knockdown only of sh*CTNNB1*, 0.2 mM of IPTG was added and for cell lines with knockdown and over-expression of β-catenin, 0.2 mM of IPTG and 1 μg/ml DOX was added to the differentiation media. For Wnt induction experiments on Y654F mutant cell line, either 100 ng/ml recombinant human Wnt3a protein (BioTechne, R&D Systems) or 3 μM CHIR-99021 (Millipore) was added to the differentiation media along with 0.2 mM IPTG and 1 μg/ml DOX.

#### Time-Lapse Imaging

Glass-bottomed 29 mm dishes with hESCs patterned colonies were placed in a 6-well plate with the bottom cut-out and placed on a custom-built stage mount. The stage mount was sealed, placed on a motorized positioning stage (Prior Scientific HLD 117) attached to a Nikon Elipse TE200 U (Nikon) inverted epifluorescent microscope and infused with a flow of mixed 5% CO_2_, 95% air (Airgas) to maintain the pH of the differentiation media. The stage, condenser and objectives were all encased in Plexiglass wherein a forced air temperature feedback control (*in* vivo Scientific) maintained the temperature of the box at 37°C. Images were captured using a 10x objective (Plan Apo NA 0.45) at specific timepoints, usually every hour during differentiation. Auto-focus on either the far-red beads or the mScarlet-NLS channel was done with the NIS Elements (Nikon) software at every timepoint, using a two-step protocol: first step finding the focus in 3 steps in a 100 μm range, then finding the best focus in 5 steps within a 30 μm range.

#### Immunofluorescence staining and imaging

For fixation, media was removed from colonies on patterned silicone gels and 4% paraformaldehyde (PFA; Sigma) was added at room temperature for 20 min.

Samples destined for β-catenin Y654 staining were briefly washed with a solution warmed to 90°C of 0.03% Triton-X 100 (Sigma), 0.4% NaCl (Fisher), referred to as “TNS” solution, and shook vigorously for 30 seconds to remove cytosolic β-catenin, but maintain junctional β-catenin, as described on ^19^. Samples were then transferred to ice and ice-cold TNS was immediately added to rapidly cool each sample. TNS was removed and replaced with 4% PFA for a 20 min fixation at room temperature.

All fixed samples were washed 3x 10 min with PBS and blocked and permeabilized at room temperature for 2h with a solution termed “IF buffer” which contains 0.1% bovine serum albumin (Fisher), 0.2% Triton-X 100 (Sigma), 0.05% Tween-20 (Sigma), 130 mM NaCl (Fisher), 13 mM Na_2_HPO_4_ (Fisher), 3.5 mM NaH_2_PO_4_ (Fisher), and 0.05% sodium azide (Sigma), supplemented with 4% goat serum (Sigma).

Primary antibodies were diluted in IF Buffer with 4% goat serum at 4°C with gentle rocking. The following day, samples were washed 3x for 10 minutes each with IF Buffer, then incubated with secondary antibodies diluted in IF Buffer with 4% goat serum for 2h at room temperature with gentle shaking. Then, samples were washed 3 × 10 minutes each with IF Buffer, then either imaged directly to obtain images of cells that had the shCTNNB1 mTurq2-NLS marker or directly incubated with 0.5 μg/ml DAPI (Invitrogen) in PBS for 5 min. DAPI incubated samples were washed 3x with PBS, then a solution of PBS and 50% glycerol (ThermoFisher) was added to the samples before imaging.

Epifluorescent images were captured on a Nikon Eclipse TE200 U (Nikon) inverted microscope with a 10x objective (Plan Apo NA 0.45) and an ORCA Flash 4.0LT CMOS camera (Hamamatsu). Confocal images were captured either using a Nikon Eclipse Ti inverted microscope (Nikon) equipped with a 60x objective, a CSU-X1 spinning disk confocal scanner (Yokogawa), and a Zyla sCMOS camera (Andor) or a Nikon CSU-W1 SoRa Spinning Disk Microscope (Nikon) equipped with a 20x (NA 0.75 – WD 1mm) or 60x (NA 1.49 – WD 0.12mm) objective, ORCA-FusionBT camera (Hamamatsu) in the Biological Imaging Development Center – UCSF.

Primary antibodies used were: anti-E-cadherin (RRID: AB_2291471, CST, 1:200), anti-Flag (RRID: AB_259529, Sigma, 1:500), anti-Slug (RRID: AB_2239535, CST, 1:400), anti-β-catenin-Y654 (RRID: AB_10623284, Sigma, 1:50), anti-β-catenin (RRID: AB_11127855, CST, rabbit, 1:200), anti-β-catenin (ECM Biosciences, mouse, 1:250), anti-phospho-Src Family (RRID: AB_10013641, CST, 1:100).

Secondary antibodies used were Alexa Fluor 488 goat anti-mouse IgG (RRID: AB_2576208, Abcam, 1:250), Alexa Fluor 647 goat anti-mouse IgG (AB_2687948, Abcam, 1:250), Alexa Fluor 488 goat anti-rabbit IgG (RRID: AB_2630356, Abcam, 1:250), Alexa Fluor 633 goat anti-rabbit IgG (AB_2920718, Abcam, 1:250).

#### Image analysis

Images were analyzed in Fiji ^69^ using custom made scripts and using python scripts.

#### Quantitative PCR (qPCR)

Total RNA was isolated from full colonies of hESCs on silicone gels or on tissue culture plastic as indicated in the main text and Figure captions using either the TRIzol (Invitrogen) manufacturer protocol or Quick-RNA Microprep Kit (Zymo Research) with no difference in yield.

RNA was used to synthesize cDNA using M-MLV Reverse Transcriptase (BioChain) and random hexamers (Applied Biosystems) as primers. qPCR was performed with technical duplicates from 10ng of RNA per reaction using PerfeCTa SYBR Green FastMix (Quantabio) on a Mastercycler RealPlex^2^ detection system (Eppendorf). All reactions for qPCR were performed using the following conditions: 95 °C for 30 s followed by 40 cycles of a three-step reaction of denaturation at 95 °C for 10 s, annealing at 65 °C for 10 s, and further annealing at 68 °C for

20 s to reduce the likelihood of non-specific products, with reads taken at the end of each 68 °C step. At the end of each reaction, melting curves were generated to validate the quality of amplified products using the following conditions: 95 °C for 15 s, 60 °C for 15 s, ramp to 95 °C in 10 min. The mean Ct values from duplicates were used to calculate the ΔCt values relative to GAPDH expression. The means of the ΔCt values from independent experiments were used to calculate mean fold change of expression using the 2^-ΔΔCt^ method and data is reported as log2 of this resulting outcome. For each gene evaluated, the ΔCt values were used to perform statistical analysis between groups. Plots of qPCR data display individual points, representing each experimental replicate relative to the mean, and mean +/- standard deviation in a log2 scale. All primers used in this study are listed in Table S1.

#### Quantification and Statistical Analysis

Quantification in this study was performed with custom Fiji ^69^ scripts or separate python scripts on Jupyter and statistical analysis tests used in this study were generated on GraphPad Prism or with a custom python script. All replicate numbers and tests chosen are reported in Figure legends, and all p-values are displayed in graphs.

In brief, nuclear images were thresholded with the StarDist algorithm^70^ plugin in Fiji ^69^ with a custom script. Other images were thresholded with the auto-threshold function in Fiji, and the method used was written on Figure legends. For graphs generated on GraphPad Prism, data was plotted in a violin plot, where dashes represent quartiles and individual datapoints are displayed, with color shading separating different experimental replicates.

The profile of circular images was created with a custom script that first averaged all available circular images for each condition, then collected the averaged pixel values at the same distance from the centroid. A mean filter of 30 pixels was used to create the line that represents the mean intensity value of pixels at the same distance from centroid. Shading indicates the 95% confidence value of values for each distance from centroid.

## Notes

### Competing Interest Statement

The authors have declared no competing interest.

## References

1. Stern, C.D. (1992). Vertebrate gastrulation. Curr. Opin. Genet. Dev. 2, 556–561. 10.1016/S0959-437X(05)80171-6.

2. Rich, A., and Glotzer, M. (2021). Small GTPases modulate intrinsic and extrinsic forces that control epithelial folding in Drosophila embryos. Small GTPases 12, 416–428. 10.1080/21541248.2021.1926879.

3. Kumburegama, S., Wijesena, N., Xu, R., and Wikramanayake, A.H. (2011). Strabismus-mediated primary archenteron invagination is uncoupled from Wnt/β-catenin-dependent endoderm cell fate specification in Nematostella vectensis (Anthozoa, Cnidaria): Implications for the evolution of gastrulation. Evodevo 2, 2. 10.1186/2041-9139-2-2.

4. Mitrossilis, D., Röper, J.-C., Le Roy, D., Driquez, B., Michel, A., Ménager, C., Shaw, G., Le Denmat, S., Ranno, L., Dumas-Bouchiat, F., et al. (2017). Mechanotransductive cascade of Myo-II-dependent mesoderm and endoderm invaginations in embryo gastrulation. Nat. Commun. 8, 13883. 10.1038/ncomms13883.

5. Rozbicki, E., Chuai, M., Karjalainen, A.I., Song, F., Sang, H.M., Martin, R., Knölker, H.-J., MacDonald, M.P., and Weijer, C.J. (2015). Myosin-II-mediated cell shape changes and cell intercalation contribute to primitive streak formation. Nat. Cell Biol. 17, 397–408. 10.1038/ncb3138.

6. Ben-Haim, N., Lu, C., Guzman-Ayala, M., Pescatore, L., Mesnard, D., Bischofberger, M., Naef, F., Robertson, E.J., and Constam, D.B. (2006). The Nodal Precursor Acting via Activin Receptors Induces Mesoderm by Maintaining a Source of Its Convertases and BMP4. Dev. Cell 11, 313–323. 10.1016/j.devcel.2006.07.005.

7. Etoc, F., Metzger, J., Ruzo, A., Kirst, C., Yoney, A., Ozair, M.Z., Brivanlou, A.H., and Siggia, E.D. (2016). A Balance between Secreted Inhibitors and Edge Sensing Controls Gastruloid Self-Organization. Dev. Cell 39, 302–315. 10.1016/j.devcel.2016.09.016.

8. Martyn, I., Brivanlou, A.H., and Siggia, E.D. (2019). A wave of WNT signalling balanced by secreted inhibitors controls primitive streak formation in micropattern colonies of human embryonic stem cells. Development. 10.1242/dev.172791.

9. Brunet, T., Bouclet, A., Ahmadi, P., Mitrossilis, D., Driquez, B., Brunet, A.-C., Henry, L., Serman, F., Béalle, G., Ménager, C., et al. (2013). Evolutionary conservation of early mesoderm specification by mechanotransduction in Bilateria. Nat. Commun. 4, 2821. 10.1038/ncomms3821.

10. Desprat, N., Supatto, W., Pouille, P.A., Beaurepaire, E., and Farge, E. (2008). Tissue Deformation Modulates Twist Expression to Determine Anterior Midgut Differentiation in Drosophila Embryos. Dev. Cell 15, 470–477. 10.1016/j.devcel.2008.07.009.

11. Muncie, J.M., Ayad, N.M.E., Lakins, J.N., Xue, X., Fu, J., and Weaver, V.M. (2020). Mechanical Tension Promotes Formation of Gastrulation-like Nodes and Patterns Mesoderm Specification in Human Embryonic Stem Cells. Dev. Cell 55, 679–694.e11. 10.1016/j.devcel.2020.10.015.

12. Nishioka, N., Inoue, K., Adachi, K., Kiyonari, H., Ota, M., Ralston, A., Yabuta, N., Hirahara, S., Stephenson, R.O., Ogonuki, N., et al. (2009). The Hippo Signaling Pathway Components Lats and Yap Pattern Tead4 Activity to Distinguish Mouse Trophectoderm from Inner Cell Mass. Dev. Cell 16, 398–410. 10.1016/j.devcel.2009.02.003.

13. van Amerongen, R., and Nusse, R. (2009). Towards an integrated view of Wnt signaling in development. Development 136, 3205–3214. 10.1242/dev.033910.

14. Huelsken, J., Vogel, R., Brinkmann, V., Erdmann, B., Birchmeier, C., and Birchmeier, W. (2000). Requirement for β-Catenin in Anterior-Posterior Axis Formation in Mice. J. Cell Biol. 148, 567–578. 10.1083/jcb.148.3.567.

15. Funa, N.S., Schachter, K.A., Lerdrup, M., Ekberg, J., Hess, K., Dietrich, N., Honoré, C., Hansen, K., and Semb, H. (2015). β-Catenin Regulates Primitive Streak Induction through Collaborative Interactions with SMAD2/SMAD3 and OCT4. Cell Stem Cell 16, 639–652. 10.1016/j.stem.2015.03.008.

16. Ledwon, J.K., Vaca, E.E., Huang, C.C., Kelsey, L.J., McGrath, J.L., Topczewski, J., Gosain, A.K., and Topczewska, J.M. (2022). Langerhans cells and SFRP2/Wnt/beta-catenin signalling control adaptation of skin epidermis to mechanical stretching. J. Cell. Mol. Med. 26, 764–775. 10.1111/jcmm.17111.

17. Przybyla, L., Lakins, J.N., and Weaver, V.M. (2016). Tissue Mechanics Orchestrate Wnt-Dependent Human Embryonic Stem Cell Differentiation. Cell Stem Cell 19, 462–475. 10.1016/j.stem.2016.06.018.

18. Shyer, A.E., Rodrigues, A.R., Schroeder, G.G., Kassianidou, E., Kumar, S., and Harland, R.M. (2017). Emergent cellular self-organization and mechanosensation initiate follicle pattern in the avian skin. Science (80-. )., 1–17. 10.1126/science.aai7868.

19. Röper, J.-C., Mitrossilis, D., Stirnemann, G., Waharte, F., Brito, I., Fernandez-Sanchez, M.-E., Baaden, M., Salamero, J., and Farge, E. (2018). The major β-catenin/E-cadherin junctional binding site is a primary molecular mechano-transductor of differentiation in vivo. Elife 7, 152–160. 10.7554/eLife.33381.

20. Nguyen, N.M., Merle, T., Broders-Bondon, F., Brunet, A.C., Battistella, A., Land, E.B.L., Sarron, F., Jha, A., Gennisson, J.L., Röttinger, E., et al. (2022). Mechano-biochemical marine stimulation of inversion, gastrulation, and endomesoderm specification in multicellular Eukaryota. Front. Cell Dev. Biol. 10, 1–21. 10.3389/fcell.2022.992371.

21. Xu, W., and Kimelman, D. (2007). Mechanistic insights from structural studies of β-catenin and its binding partners. J. Cell Sci. 120, 3337–3344. 10.1242/jcs.013771.

22. Roura, S., Miravet, S., Piedra, J., García De Herreros, A., and Duñachl, M. (1999). Regulation of E-cadherin/catenin association by tyrosine phosphorylation. J. Biol. Chem. 274, 36734–36740. 10.1074/jbc.274.51.36734.

23. Yamaguchi, T.P., Takada, S., Yoshikawa, Y., Wu, N., and McMahon, A.P. (1999). T (Brachyury) is a direct target of Wnt3a during paraxial mesoderm specification. Genes Dev. 13, 3185–3190. 10.1101/gad.13.24.3185.

24. Arnold, S.J., Stappert, J., Bauer, A., Kispert, A., Herrmann, B.G., and Kemler, R. (2000). Brachyury is a target gene of the Wnt/β-catenin signaling pathway. Mech. Dev. 91, 249–258. 10.1016/S0925-4773(99)00309-3.

25. Miroshnikova, Y.A., Mouw, J.K., Barnes, J.M., Pickup, M.W., Lakins, J.N., Kim, Y., Lobo, K., Persson, A.I., Reis, G.F., McKnight, T.R., et al. (2016). Tissue mechanics promote IDH1-dependent HIF1α–tenascin C feedback to regulate glioblastoma aggression. Nat. Cell Biol. 18, 1336–1345. 10.1038/ncb3429.

26. Kowarz, E., Löscher, D., and Marschalek, R. (2015). Optimized Sleeping Beauty transposons rapidly generate stable transgenic cell lines. Biotechnol. J. 10, 647–653. 10.1002/biot.201400821.

27. Heasman, J. (1994). Overexpression of cadherins and underexpression of ?-catenin inhibit dorsal mesoderm induction in early Xenopus embryos. Cell 79, 791–803. 10.1016/0092-8674(94)90069-8.

28. Funayama, N., Fagotto, F., McCrea, P., and Gumbiner, B.M. (1995). Embryonic axis induction by the armadillo repeat domain of beta-catenin: evidence for intracellular signaling. J. Cell Biol. 128, 959–968. 10.1083/jcb.128.5.959.

29. Ou, G., Thakar, D., Tung, J.C., Miroshnikova, Y.A., Dufort, C.C., Gutierrez, E., Groisman, A., and Weaver, V.M. (2016). Visualizing mechanical modulation of nanoscale organization of cell-matrix adhesions. Integr. Biol. 8, 795–804. 10.1039/C6IB00031B.

30. Muncie, J.M., Falcón-Banchs, R., Lakins, J.N., Sohn, L.L., and Weaver, V.M. (2019). Patterning the Geometry of Human Embryonic Stem Cell Colonies on Compliant Substrates to Control Tissue-Level Mechanics. J. Vis. Exp. 10.3791/60334.

31. Winnier, G., Blessing, M., Labosky, P.A., and Hogan, B.L.M. (1995). Bone morphogenetic protein-4 is required for mesoderm formation and patterning in the mouse. Genes Dev. 9, 2105–2116. 10.1101/gad.9.17.2105.

32. Shi, Y., and Massagué, J. (2003). Mechanisms of TGF-β Signaling from Cell Membrane to the Nucleus. Cell 113, 685–700. 10.1016/S0092-8674(03)00432-X.

33. Warmflash, A., Sorre, B., Etoc, F., Siggia, E.D., and Brivanlou, A.H. (2014). A method to recapitulate early embryonic spatial patterning in human embryonic stem cells. Nat. Methods 11, 847–854. 10.1038/nMeth.3016.

34. Galceran, J., Hsu, S.-C., and Grosschedl, R. (2001). Rescue of a Wnt mutation by an activated form of LEF-1: Regulation of maintenance but not initiation of Brachyury expression. Proc. Natl. Acad. Sci. 98, 8668–8673. 10.1073/pnas.151258098.

35. Fuentealba, L.C., Eivers, E., Ikeda, A., Hurtado, C., Kuroda, H., Pera, E.M., and De Robertis, E.M. (2007). Integrating Patterning Signals: Wnt/GSK3 Regulates the Duration of the BMP/Smad1 Signal. Cell 131, 980–993. 10.1016/j.cell.2007.09.027.

36. Webb, D.J., Donais, K., Whitmore, L.A., Thomas, S.M., Turner, C.E., Parsons, J.T., and Horwitz, A.F. (2004). FAK–Src signalling through paxillin, ERK and MLCK regulates adhesion disassembly. Nat. Cell Biol. 6, 154–161. 10.1038/ncb1094.

37. Gayrard, C., Bernaudin, C., Déjardin, T., Seiler, C., and Borghi, N. (2018). Src- and confinement-dependent FAK activation causes E-cadherin relaxation and β-catenin activity. J. Cell Biol. 217, 1063–1077. 10.1083/jcb.201706013.

38. Gayrard, C., Bernaudin, C., Déjardin, T., Seiler, C., and Borghi, N. (2018). Src- and confinement-dependent FAK activation causes E-cadherin relaxation and β-catenin activity. J. Cell Biol. 217, 1063–1077. 10.1083/jcb.201706013.

39. Shi, J., Severson, C., Yang, J., Wedlich, D., and Klymkowsky, M.W. (2011). Snail2 controls mesodermal BMP/Wnt induction of neural crest. Development 138, 3135–3145. 10.1242/dev.064394.

40. González-Sancho, J.M., Aguilera, O., García, J.M., Pendás-Franco, N., Peña, C., Cal, S., de Herreros, A.G., Bonilla, F., and Muñoz, A. (2005). The Wnt antagonist DICKKOPF-1 gene is a downstream target of β-catenin/TCF and is downregulated in human colon cancer. Oncogene 24, 1098–1103. 10.1038/sj.onc.1208303.

41. Gomez, E.W., Chen, Q.K., Gjorevski, N., and Nelson, C.M. (2010). Tissue geometry patterns epithelial-mesenchymal transition via intercellular mechanotransduction. J. Cell. Biochem., n/a-n/a. 10.1002/jcb.22545.

42. Van Veelen, W., Le, N.H., Helvensteijn, W., Blonden, L., Theeuwes, M., Bakker, E.R.M., Franken, P.F., Van Gurp, L., Meijlink, F., Van Der Valk, M.A., et al. (2011). β-catenin tyrosine 654 phosphorylation increases Wnt signalling and intestinal tumorigenesis. Gut 60, 1204–1212. 10.1136/gut.2010.233460.

43. Tucci, V., Kleefstra, T., Hardy, A., Heise, I., Maggi, S., Willemsen, M.H., Hilton, H., Esapa, C., Simon, M., Buenavista, M.T., et al. (2014). Dominant β-catenin mutations cause intellectual disability with recognizable syndromic features. J. Clin. Invest. 124, 1468–1482. 10.1172/JCI70372.

44. Huang, C.F., Gottardi, C.J., and Mrksich, M. (2022). Tyrosine phosphatase activity is restricted by basic charge substituting mutation of substrates. Sci. Rep. 12, 1–9. 10.1038/s41598-022-19133-4.

45. Fernández-Sánchez, M.E., Barbier, S., Whitehead, J., Béalle, G., Michel, A., Latorre-Ossa, H., Rey, C., Fouassier, L., Claperon, A., Brullé, L., et al. (2015). Mechanical induction of the tumorigenic β-catenin pathway by tumour growth pressure. Nature 523, 92–95. 10.1038/nature14329.

46. Valenta, T., Hausmann, G., and Basler, K. (2012). The many faces and functions of β- catenin. EMBO J. 31, 2714–2736. 10.1038/emboj.2012.150.

47. Orsulic, S., Huber, O., Aberle, H., Arnold, S., and Kemler, R. (1999). E-cadherin binding prevents β-catenin nuclear localization and β-catenin/LEF-1-mediated transactivation. J. Cell Sci. 112, 1237–1245. 10.1242/jcs.112.8.1237.

48. van der Wal, T., and van Amerongen, R. (2020). Walking the tight wire between cell adhesion and WNT signalling: a balancing act for β-catenin. Open Biol. 10. 10.1098/rsob.200267.

49. Cook, D., Fry, M.J., Hughes, K., Sumathipala, R., Woodgett, J.R., and Dale, T.C. (1996). Wingless inactivates glycogen synthase kinase-3 via an intracellular signalling pathway which involves a protein kinase C. EMBO J. 15, 4526–4536. 8887544/.

50. Cselenyi, C.S., Jernigan, K.K., Tahinci, E., Thorne, C.A., Lee, L.A., and Lee, E. (2008). LRP6 transduces a canonical Wnt signal independently of Axin degradation by inhibiting GSK3’s phosphorylation of β-catenin. Proc. Natl. Acad. Sci. 105, 8032–8037. 10.1073/pnas.0803025105.

51. McCrea, P.D., Maher, M.T., and Gottardi, C.J. (2015). Nuclear Signaling from Cadherin Adhesion Complexes. In, pp. 129–196. 10.1016/bs.ctdb.2014.11.018.

52. Anzalone, A. V., Randolph, P.B., Davis, J.R., Sousa, A.A., Koblan, L.W., Levy, J.M., Chen, P.J., Wilson, C., Newby, G.A., Raguram, A., et al. (2019). Search-and-replace genome editing without double-strand breaks or donor DNA. Nature 576, 149–157. 10.1038/s41586-019-1711-4.

53. Chhabra, S., Liu, L., Goh, R., Kong, X., and Warmflash, A. (2019). Dissecting the dynamics of signaling events in the BMP, WNT, and NODAL cascade during self-organized fate patterning in human gastruloids. PLOS Biol. 17, e3000498. 10.1371/journal.pbio.3000498.

54. Arnold, S.J., and Robertson, E.J. (2009). Making a commitment: cell lineage allocation and axis patterning in the early mouse embryo. Nat. Rev. Mol. Cell Biol. 10, 91–103. 10.1038/nrm2618.

55. Bailles, A., Collinet, C., Philippe, J.-M., Lenne, P.-F., Munro, E., and Lecuit, T. (2019). Genetic induction and mechanochemical propagation of a morphogenetic wave. Nature 572, 467–473. 10.1038/s41586-019-1492-9.

56. Tyser, R.C.V., and Srinivas, S. (2022). Recent advances in understanding cell types during human gastrulation. Semin. Cell Dev. Biol. 131, 35–43. 10.1016/j.semcdb.2022.05.004.

57. Weatherbee, B.A.T., Gantner, C.W., Iwamoto-Stohl, L.K., Daza, R.M., Hamazaki, N., Shendure, J., and Zernicka-Goetz, M. (2023). A model of the post-implantation human embryo derived from pluripotent stem cells. Nature. 10.1038/s41586-023-06368-y.

58. Oldak, B., Wildschutz, E., Bondarenko, V., Aguilera-Castrejon, A., Zhao, C., Tarazi, S., Comar, M.-Y., Ashouokhi, S., Lokshtanov, D., Roncato, F., et al. (2023). Transgene-Free Ex Utero Derivation of A Human Post-Implantation Embryo Model Solely from Genetically Unmodified Naïve PSCs. bioRxiv, 2023.06.14.544922. 10.1101/2023.06.14.544922.

59. Karzbrun, E., Khankhel, A.H., Megale, H.C., Glasauer, S.M.K., Wyle, Y., Britton, G., Warmflash, A., Kosik, K.S., Siggia, E.D., Shraiman, B.I., et al. (2021). Human neural tube morphogenesis in vitro by geometric constraints. Nature 599, 268–272. 10.1038/s41586-021-04026-9.

60. Veenvliet, J. V., Bolondi, A., Kretzmer, H., Haut, L., Scholze-Wittler, M., Schifferl, D., Koch, F., Guignard, L., Kumar, A.S., Pustet, M., et al. (2020). Mouse embryonic stem cells self-organize into trunk-like structures with neural tube and somites. Science (80-. ). 370. 10.1126/science.aba4937.

61. van den Brink, S.C., Alemany, A., van Batenburg, V., Moris, N., Blotenburg, M., Vivié, J., Baillie-Johnson, P., Nichols, J., Sonnen, K.F., Martinez Arias, A., et al. (2020). Single-cell and spatial transcriptomics reveal somitogenesis in gastruloids. Nature 582, 405–409. 10.1038/s41586-020-2024-3.

62. Cederquist, G.Y., Asciolla, J.J., Tchieu, J., Walsh, R.M., Cornacchia, D., Resh, M.D., and Studer, L. (2019). Specification of positional identity in forebrain organoids. Nat. Biotechnol. 37, 436–444. 10.1038/s41587-019-0085-3.

63. Sozen, B., Conkar, D., and Veenvliet, J. V. (2022). Carnegie in 4D? Stem-cell-based models of human embryo development. Semin. Cell Dev. Biol. 131, 44–57. 10.1016/j.semcdb.2022.05.023.

64. Przybyla, L., Lakins, J.N., Sunyer, R., Trepat, X., and Weaver, V.M. (2016). Monitoring developmental force distributions in reconstituted embryonic epithelia. Methods 94, 101–113. 10.1016/j.ymeth.2015.09.003.

65. Xue, X., Sun, Y., Resto-Irizarry, A.M., Yuan, Y., Aw Yong, K.M., Zheng, Y., Weng, S., Shao, Y., Chai, Y., Studer, L., et al. (2018). Mechanics-guided embryonic patterning of neuroectoderm tissue from human pluripotent stem cells. Nat. Mater. 17, 633–641. 10.1038/s41563-018-0082-9.

66. Mátés, L., Chuah, M.K.L., Belay, E., Jerchow, B., Manoj, N., Acosta-Sanchez, A., Grzela, D.P., Schmitt, A., Becker, K., Matrai, J., et al. (2009). Molecular evolution of a novel hyperactive Sleeping Beauty transposase enables robust stable gene transfer in vertebrates. Nat. Genet. 41, 753–761. 10.1038/ng.343.

67. Klein, S.M., Manoharan, V.N., Pine, D.J., and Lange, F.F. (2003). Preparation of monodisperse PMMA microspheres in nonpolar solvents by dispersion polymerization with a macromonomeric stabilizer. Colloid Polym. Sci. 282, 7–13. 10.1007/s00396-003-0915-0.

68. Yoshie, H., Koushki, N., Kaviani, R., Tabatabaei, M., Rajendran, K., Dang, Q., Husain, A., Yao, S., Li, C., Sullivan, J.K., et al. (2018). Traction Force Screening Enabled by Compliant PDMS Elastomers. Biophys. J. 114, 2194–2199. 10.1016/j.bpj.2018.02.045.

69. Schindelin, J., Arganda-Carreras, I., Frise, E., Kaynig, V., Longair, M., Pietzsch, T., Preibisch, S., Rueden, C., Saalfeld, S., Schmid, B., et al. (2012). Fiji: an open-source platform for biological-image analysis. Nat. Methods 9, 676–682. 10.1038/nmeth.2019.

70. Schmidt, U., Weigert, M., Broaddus, C., and Myers, G. (2018). Cell Detection with Star-Convex Polygons. In, pp. 265–273. 10.1007/978-3-030-00934-2_30.

